# Cryo-EM structures of SARS-CoV-2 Omicron BA.2 spike

**DOI:** 10.1101/2022.04.07.487528

**Authors:** Victoria Stalls, Jared Lindenberger, Sophie M-C. Gobeil, Rory Henderson, Rob Parks, Maggie Barr, Margaret Deyton, Mitchell Martin, Katarzyna Janowska, Xiao Huang, Aaron May, Micah Speakman, Esther Beaudoin, Bryan Kraft, Xiaozhi Lu, Robert J Edwards, Amanda Eaton, David C. Montefiori, Wilton Williams, Kevin O. Saunders, Kevin Wiehe, Barton F. Haynes, Priyamvada Acharya

**Author notes:** Correspondence (B.F.H.), (P.A.).

## Abstract

The BA.2 sub-lineage of the SARS-CoV-2 Omicron variant has gained in proportion relative to BA.1. As differences in spike (S) proteins may underlie differences in their pathobiology, here we determine cryo-EM structures of a BA.2 S ectodomain and compare these to previously determined BA.1 S structures. BA.2 Receptor Binding Domain (RBD) mutations induced remodeling of the internal RBD structure resulting in its improved thermostability and tighter packing within the 3-RBD-down spike. In the S2 subunit, the fusion peptide in BA.2 was less accessible to antibodies than in BA.1. Pseudovirus neutralization and spike binding assays revealed extensive immune evasion while defining epitopes of two RBD-directed antibodies, DH1044 and DH1193, that bound the outer RBD face to neutralize both BA.1 and BA.2. Taken together, our results indicate that stabilization of the 3-RBD-down state through interprotomer RBD-RBD packing is a hallmark of the Omicron variant, and reveal differences in key functional regions in the BA.1 and BA.2 S proteins.

## Introduction

The SARS-CoV-2 Omicron B.1.1.529 (or Nexstrain 21M) variant, first detected in November 2021, includes several sub-lineages, including BA.1 (B.1.1.529.1 or Nextstrain clade 21K), BA.2 (B.1.1.529.2 or Nextstrain clade 21L) and BA.3 (B.1.1.529.3 or Nextstrain clade 21M) (**Figure 1 and S1**) (Hadfield et al., 2018, Sagulenko et al., 2018). BA.1 was the first of the Omicron sub-lineages to rapidly spread worldwide, although over the past weeks the proportion of reported BA.2 sequences has increased relative to BA.1, and BA.2 has now overtaken BA.1 to become the dominant coronavirus variant in the US (https://covid.cdc.gov/covid-data-tracker/#variant-proportions) (Viana et al., 2022). The Omicron variant is characterized by its high number of mutations in the spike (S) protein. BA.1 and BA.2 have 20 S protein mutations in common (relative to the D614G spike), although they each have 13 and 8 unique mutations, respectively. These differences may be responsible for differences in S protein mediated properties such as host cell entry, viral transmission, and immune recognition.

**Figure 1.**
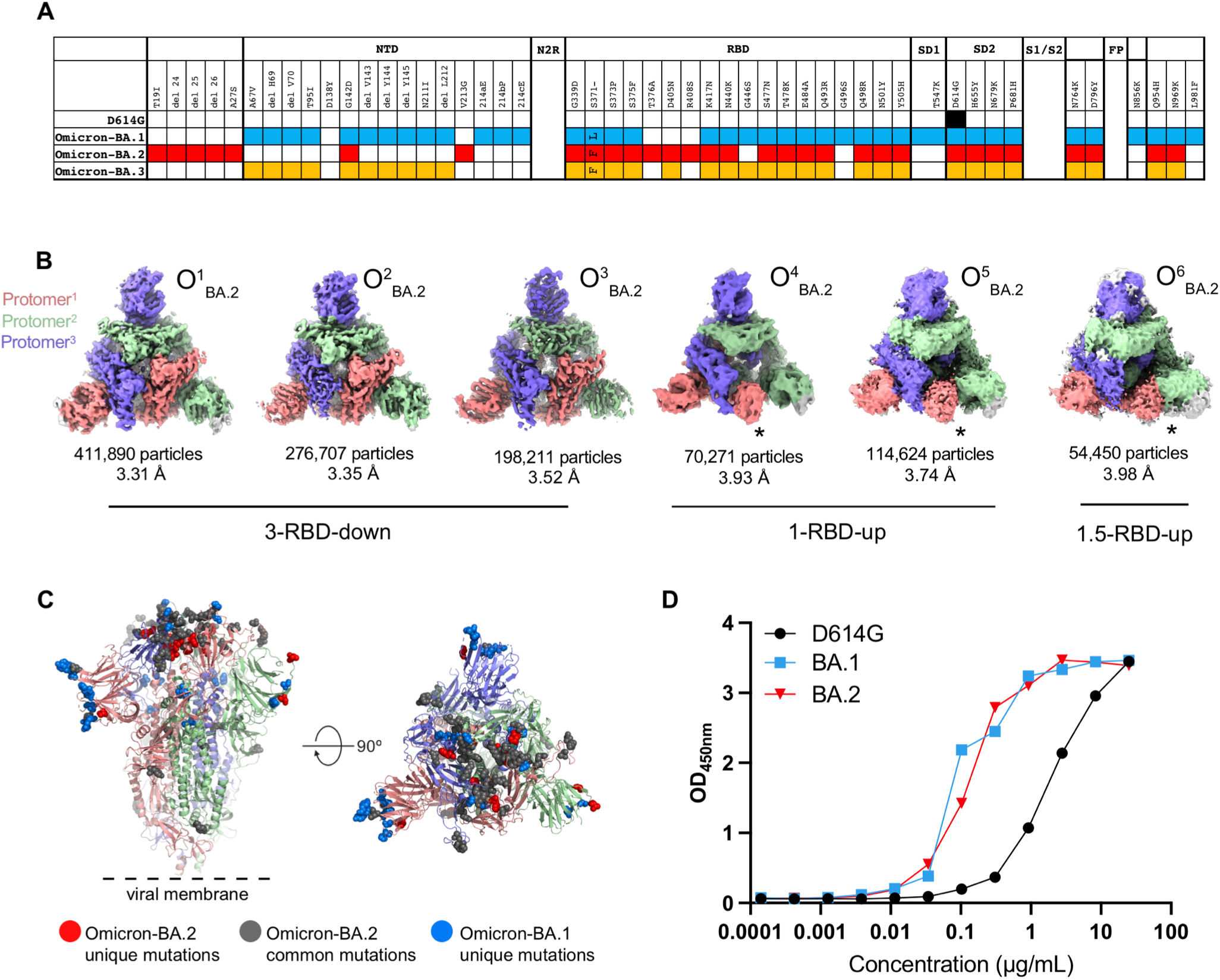
Structural characterization of SARS-CoV-2 Omicron-BA.2 spike (S) protein, Related to Figures S1-S4, Table 1. **A**. Comparison of residue changes in the S ectodomain (S-GSAS) of SARS-CoV-2 D614G and Omicron variant sub-lineages. Residue changes from the original Wuhan strain are color coded for the variants: D614G (black), BA.1 (blue), BA.2 (red), and BA.3 (yellow). **B**. Cryo-EM reconstructions of Omicron-BA.2 S protein 3-RBD-down (O^1^_BA.2_: EMD-26433, PDB 7UB0; O^2^_BA.2_: EMD-26435, PDB 7UB5; O^3^_BA.2_: EMD-26436, PDB 7UB6), 1-RBD-up (O^4^_BA.2_ : EMD-26644, O^5^_BA.2_ : EMD-26647), and 1.5-RBD-up (O^6^_BA.2_ : EMD-26643) states, colored by protomer, and viewed from the host cell membrane. In the RBD-up reconstructions, the “up” RBD is indicated by an asterisk (*). **C**. Omicron-BA.2 S 3-RBD-down structure (O^1^_BA.2_: EMD-26433, PDB 7UB0) colored by protomer, with common mutations shown as gray spheres, BA.2 unique mutations colored red, and BA.1 unique mutations colored blue. **D**. ACE-2 binding to SARS-CoV-2 S proteins measured by ELISA.

The BA.1 and BA.2 S proteins differ substantially in their N terminal domains (NTDs) with only the G142D substitution shared between the two (https://www.gisaid.org/hcov19-variants/) (**Figure 1A**). The G142D substitution also occurred in Delta VOC sub-lineages and has been associated with immune evasion and high viral loads (Shen et al., 2021). Notably, the BA.2 S protein NTD lacks the H69-V70 deletion (ΔH69-V70) that is present in BA.1, as well as in the Alpha (B.1.1.7) and a mink-associated (ΔFV) variant (Gobeil et al., 2021b, Meng et al., 2021). The BA.2 NTD also lacks the deletion of residues 143-145, and insertion of 3 residues at position 214 (EPE). The Receptor Binding Domains (RBDs) of BA.1 and BA.2 are more similar with 12 shared mutations, including two, S373P and S375F, that occur in an RBD loop previously implicated in mediating RBD-RBD packing in the 3-RBD-down BA.1 S protein (Gobeil et al., 2022). Residue S371, also part of this interfacial RBD loop, is mutated to Leu in BA.1 or to Phe in BA.2. The Omicron BA.2 S protein harbors an additional amino acid substitution, T376A, within this interfacial loop. RBD mutations that occur in the BA.2 S protein but not in BA.1 are T376A, D405N and R408S, whereas G446S and G496S occur in BA.1 but not in the BA.2 S protein. The BA.2 S protein lacks the SD1 T457K, and S2 N856K and L981F substitutions that occur in BA.1. All other mutations outside the RBD/NTD region are conserved between the two (**Figure 1A**).

We and others have described structures of the Omicron BA.1 spike (Zhou, 2021, Mannar et al., 2022, Cerutti et al., 2022, Cui et al., 2022, McCallum et al., 2022, Ye et al., 2022, Gobeil et al., 2022). To understand the differences between the BA.1 and BA.2 S proteins, here we determine cryo-EM structures of the BA.2 S protein ectodomain. The BA.2 S cryo-EM dataset was dominated by 3-RBD-down populations, although we also resolved RBD “up” populations. The dominance of the 3-RBD-down state was driven by improved RBD-RBD packing. An RBD interfacial loop, containing the S373P and S375F residue substitutions, that we had previously identified in the BA.1 S protein as a facilitator of RBD-RBD packing in the 3-RBD-down state (Gobeil et al., 2022), incorporates additional residue substitutions in the BA.2 S protein that drove even closer interactions between the RBDs in the 3-RBD-down structures. These additional mutations also bolstered internal packing within each RBD resulting in a more stable fold relative to the BA.1 RBD. We found differences in the S2 subunit between the BA.1 and BA.2 S proteins, including reduced accessibility of the fusion peptide (FP) in BA.2 to FP-directed antibodies. While we observed extensive immune evasion, two antibodies DH1044 and DH1193, that bind the outer face of the RBD neutralized BA.1 and BA.2 in pseudovirus neutralization assays. Taken together, our results demonstrate key differences in the Omicron BA.2 S protein architecture compared to BA.1 that may drive the differences in their biological properties.

## Results

### Structural diversity and ACE2 binding of the SARS-CoV-2 Omicron BA.2 S protein

We determined cryo-EM structures of the BA.2 S ectodomain using our previously described S-GSAS-D614G platform (**Figures 1, S2-S6, Table 1**) (Gobeil et al., 2021a, Gobeil et al., 2021b). We identified 3-RBD-down (O^1^_BA.2_, O^2^_BA.2_ and O^3^_BA.2_) and RBD-up (O^4^_BA.2_, O^5^_BA.2_ and O^6^_BA.2_) states in a roughly 3:1 ratio (**Figure 1B**), which was higher than the ∼2:1 ratio we had observed for the BA.1 spike (Gobeil et al., 2022). The 1-RBD-up populations, O^4^_BA.2_ and O^5^_BA.2_, differed primarily by the position of the up-RBD, while O^6^_BA.2_ had 1 RBD in the “up” position and a second RBD was partially up. Of the three BA.2 3-RBD-down S protein structures O^2^_BA.2_ and O^3^_BA.2_ were more symmetric than the third, O^1^_BA.2_, which showed asymmetry in RBD dispositions between its three protomers (**Figure S7**), with O^2^_BA.2_ and O^3^_BA.2_ being more similar to each other than either was to the O^1^_BA.2_ structure (**Figure S8**). Despite its heavily mutated RBD (**Figure 1C**), the Omicron BA.2 S ectodomain showed robust binding to an ACE2 receptor ectodomain construct, at levels similar to that of the BA.1 S ectodomain and higher than the D614G spike, showing that the large number of RBD mutations and the increased propensity for the 3-RBD-down state does not impair the ability of the Omicron BA.2 spike to bind ACE2 under the conditions of the ELISA assay (**Figure 1D**).

**Table 1:** Cryo-EM data collection and refinements statistics S-GSAS-Omicron-BA.2, related to Figure 1.

|  | 3-down |  |  | 1-up |  | 1.5-up |
| --- | --- | --- | --- | --- | --- | --- |
| **PDB ID** | 7UB0 | 7UB5 | 7UB6 | n/a | n/a | n/a |
| **EMDB ID** | 26433 | 26435 | 26436 | 26644 | 26647 | 26643 |
| **Data Collection and processing** |  |  |  |  |  |  |
| Microscope |  |  |  | FEI Titan Krios |  |  |
| Detector |  |  |  | Gatan K3 |  |  |
| Magnification |  |  |  | 81000 |  |  |
| Voltage (kV) |  |  |  | 300 |  |  |
| Electron exposure (e-/Å<sup>2</sup>) |  |  |  | 54.7 |  |  |
| Defocus Range (μm) |  |  |  | 2.6 to 0.8 |  |  |
| Pixel size (Å) |  |  |  | 1.08 |  |  |
| Reconstruction software |  |  |  | cryoSPARC |  |  |
| Symmetry imposed | C1 | C1 | C1 | C1 | C1 | C1 |
| Initial particle images (no.) |  |  |  |  |  |  |
| Final particle images (no.) | 411,890 | 276,707 | 198,211 | 70,271 | 114,624 | 54,450 |
| Map resolution (Å) | 3.31 | 3.35 | 3.52 | 3.93 | 3.74 | 3.98 |
| FSC threshold | 0.143 | 0.143 | 0.143 | 0.143 | 0.143 | 0.143 |
| **Refinement** |  |  |  |  |  |  |
| Initial model used | 7KDK | 7KDK | 7KDK |  |  |  |
| **Model composition** |  |  |  |  |  |  |
| Nonhydrogen atoms | 25,039 | 24,963 | 24,762 |  |  |  |
| Protein residues | 3,112 | 3,107 | 3,105 |  |  |  |
| **R.M.S. deviations** |  |  |  |  |  |  |
| Bond lengths (Å) | 0.004 | 0.004 | 0.004 |  |  |  |
| Bond angles (°) | 0.651 | 0.663 | 0.669 |  |  |  |
| **Validation** |  |  |  |  |  |  |
| MolProbity score | 1.22 | 1.29 | 1.24 |  |  |  |
| Clashscore | 3.25 | 2.74 | 2.96 |  |  |  |
| Poor rotamers (%) | 0 | 0 | 0 |  |  |  |
| EM ringer score | 3.44 | 3.13 | 2.97 |  |  |  |
| **Ramachandran plot** |  |  |  |  |  |  |
| Favored regions (%) | 97.49 | 96.47 | 97.19 |  |  |  |
| Allowed (%) | 2.51 | 3.53 | 2.81 |  |  |  |
| Disallowed regions (%) | 0 | 0 | 0 |  |  |  |

### Thermostability of the SARS-CoV-2 Omicron BA.2 spike and RBD

We tested the thermostability of the D614G, BA.1 and BA.2 S protein ectodomains and of the corresponding monomeric RBD constructs using a differential scanning fluorimetry (DSF) assay that measures changes in intrinsic fluorescence of a protein as a function of temperature (**Figures 2, S2 and S9**). The WT RBD showed an inflection temperature (T_i_) of ∼54.4 °C (**Figure 2B**). The BA.1 RBD was substantially less stable with a T_i_ of ∼47.7 °C. The reduced stability of the BA.1 RBD relative to the WT RBD is in close agreement with published reports (Lin et al., 2022). The thermostability of the BA.2 RBD, with a T_i_ of ∼50.4 °C, was intermediate between that of the WT and BA.1 RBD. The D614G, BA.1 and BA.2 S proteins showed characteristic DSF profiles (Edwards et al., 2021). We have previously shown that the DSF profiles of SARS-CoV-2 S ectodomain, particularly the first inflection temperature (T_i_ #1) is sensitive to spike stability. We observed a substantial decrease of T_i_ #1 for the BA.1 S ectodomain but not for BA.2 S (**Figures 2A** and **S2**). Taken together our results show stabilization of both the monomeric RBD (or RBD-only) construct and the S ectodomain in BA.2 relative to BA.1.

**Figure 2.**
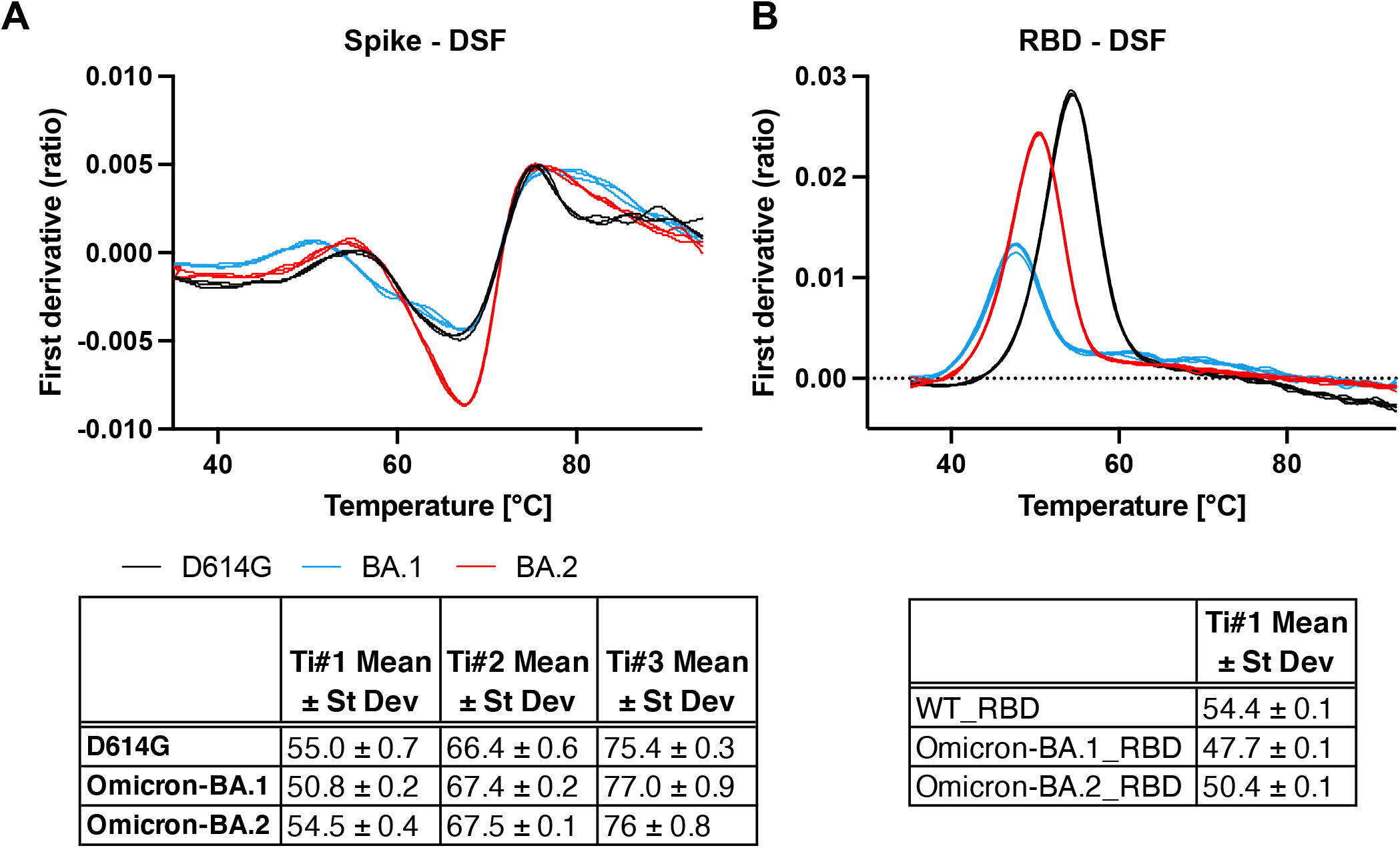
Thermostability of SARS-CoV-2 S ectodomain and RBD, Related to Figures S2 and S9. **A**. Top. DSF profiles of the SARS-CoV-2 S ectodomains showing changes in protein intrinsic fluorescence (expressed as a ratio between fluorescence at 350 and 330 nm) with temperature. For each S protein construct, three overlaid curves (technical replicates) are shown. Bottom. Maxima and minima indicate inflection temperatures, *T*_i_ #1-3 are represented as mean ± standard deviation from three technical replicates. **B**. Same as A. but for a monomeric RBD construct.

### Interprotomer RBD packing in the 3-RBD-down Omicron BA.2 S protein

Cryo-EM maps of the 3-RBD-down (or closed) SARS-CoV-2 S protein have typically exhibited considerable disorder in the RBD densities, indicative of high mobility (Gobeil et al., 2021b). The most notably visible feature of the Omicron BA.2 S protein 3-RBD-down structures was their tightly packed and well-resolved RBDs (**Figure 1B**). We had observed close interprotomer RBD packing in the 3-RBD-down structures of the Omicron BA.1 S protein (Gobeil et al., 2022); this appeared further reinforced in the BA.2 spike with the RBDs packed closer together (**Figures 1B, 1C, 3, S5** and **S10**). In the BA.1 spike, RBD-RBD contacts were mediated by a triad of acquired amino acid substitutions S371L/S373P/S375F that occurred within an interfacial RBD loop, with the S373P substitution restructuring the loop (relative to its structure in the D614G spike) facilitating close interaction with another RBD interfacial loop harboring the Y505H substitution in the adjacent protomer (**Figures 3D-F** and **S10**) (Gobeil et al., 2022). The S373P, S375F and the Y505H substitutions observed at the RBD-RBD interface in the 3-RBD-down BA.1 S protein are also present in BA.2 (**Figures 1A, 3A-C and S10**). BA.2 has a S371F substitution instead of S371L in BA.1 (**Figures 1A, 3B and S10**) and an additional T376A substitution in this interfacial RBD loop (**Figures 1A, 3C, 3F** and **S10**). The BA.2 S371F substitution results in van der Waals interaction of the bulkier F371 side chain with F342, resulting in closer packing of the two helical stretches 339-342 and 367-371 within the RBD (**Figure 3B**). Due to the packing of the F371 side chain against the 367-371 helical turn within the BA.2 RBD, the region between residues 371 and 373 flips its position, bringing residue P373 closer to the RBD of the adjacent protomer allowing its H505 side chain to stack against P373 (**Figure 3B**). Additionally, due to the closer proximity of these two adjacent RBD interfacial loops, the H505 side chain can form an interprotomer hydrogen bond with the main chain carbonyl of residue A372. The loop bearing the Y505H substitution also incorporates two additional VOC mutations, N501Y and Q498R, that occur in both BA.1 and BA.2. Y501 and R498 engage in an intra-loop cation-π interaction conferring a defined structure to this region that remains invariant between BA.1 and BA.2 (**Figures 3B and 3E**). Although the internal structure of this cation-π stabilized loop remains invariant, as a consequence of the closer inter-protomer interaction in BA.2 involving residue H505, this entire region spanning residues 494-507 is pulled closer to the adjacent protomer. Resulting from the close interprotomer packing of the two interfacial RBD loops, the side chain of K440 (from the N440K substitution that occurs in both BA.1 and BA.2) is positioned to engage in a hydrogen bond with the main chain of residue T500 from the adjacent protomer RBD (**Figures 3B** and **S10**).

**Figure 3.**
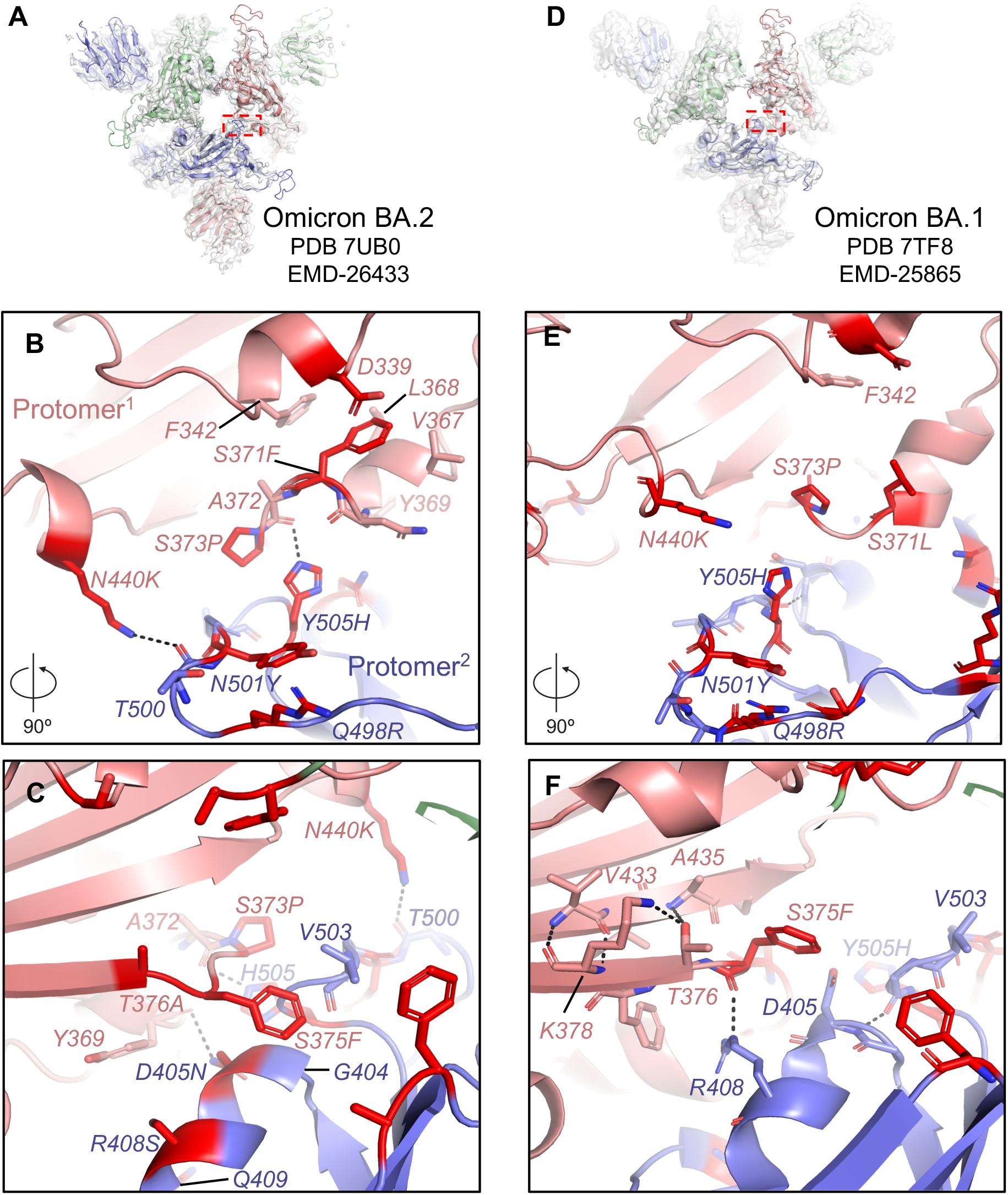
Omicron BA.2 S mutations induce RBD interfacial loop remodeling facilitating tight packing of the 3-RBD-down state, Related to Figures S5, S7, S8, and S10. **A**. View of the Omicron BA.2 S protein O^1^_BA.2_ state with the red dotted rectangle indicating the region shown in figures **B** and **C. D**. Same as **A** but for the Omicron BA.1 S protein O1 state (O^1^_BA.1_) with the red rectangle indicating the region shown in figures **E** and **F. B** and **C**, 90° rotated views of the interface between two RBDs in the 3-RBD-down O^1^_BA.2_ structure. The sites of mutation are colored red and the residues shown in sticks. **E** and **F**. Same as **B** and **C** but for the Omicron BA.1 S protein O^1^_BA.1_ structure.

Another structural change in this key 371-376 interfacial loop is orchestrated by the T376A substitution in the BA.2 RBD (**Figures 3C** and **S10**). In the BA.1 RBD, residue T376 is part of a β-strand and its side chain engages in hydrogen bond interactions with the main chain of A435 in the adjacent β-strand (**Figure 3F**). The loss of the side chain hydroxyl due to the T376A substitution in BA.2 disrupts this hydrogen bond allowing A376 to move away from the β-sheet, and as it does so, it pulls along the region around the S375F substitution so that F375 can now reach over to stack against the G404-Q409 helix and the V503 side chain of the adjacent protomer (**Figure 3C** and **S10**). Together with the restructuring of the 371-376 loop, concerted changes also occur in the adjacent protomer RBD that are key for forming the new interactions. The BA.2 S R408S substitution results in the disruption of an interprotomer hydrogen bond that the residue R408 side chain makes with the main chain carbonyl of residue 375 (**Figure**s **3C, 3F** and **S10**). This releases F375 and facilitates its movement towards the adjacent RBD. The loss of the interprotomer H-bond due to the R408S substitution may be compensated by the BA.2 S D405N substitution that mediates an interprotomer H-bond with the main chain carbonyl of Y369 (**Figure 3C**).

Taken together our results provide evidence for enhanced interprotomer RBD-RBD packing in the 3-RBD-down BA.2 spike relative to the BA.1 spike, orchestrated by residue substitutions that re-model interfacial RBD loops to engineer their close packing within each RBD, as well as between RBDs in the closed spike.

### Intra- and inter-protomer communication in the Omicron BA.2 3-RBD-down spike

We previously recognized a stretch of residues that connected the NTD and RBD within a protomer to be a modulator of RBD up/down transitions (**Figure 4**) (Gobeil et al., 2021a, Gobeil et al., 2022). In an RBD-down protomer, this NTD-to-RBD (“N2R”) linker interacts with the SD1 and SD2 subunits by contributing a β-strand to each subdomain. An Omicron BA.1 S protein 3-RBD-down structure (named O^1^_BA.1_; PDB 7TF8) stabilized a rearrangement in the N2R linker, thus possibly predisposing this protomer to adopt the RBD-up configuration (**Figure 4B**) (Gobeil et al., 2022). We had also found this N2R rearranged state in other variants, albeit to lesser extents, suggesting that this N2R rearranged state may be an intermediate in the RBD up/down transition. In the Omicron BA.1 S cryo-EM dataset, we had also identified another 3-RBD-down population (named O^2^_BA.1_; PDB 7TL1) that did not show this N2R rearrangement (**Figure 4B**) (Gobeil et al., 2022). Examining the N2R region in the Omicron BA.2 S 3-RBD-down structures, we found that none of them showed the N2R rearrangement that we had observed in the Omicron BA.1 O^1^_BA.1_ structure (**Figure 4A and 4B**), with the three protomers within each structure aligning well in the N2R region (**Figure 4A**).

**Figure 4.**
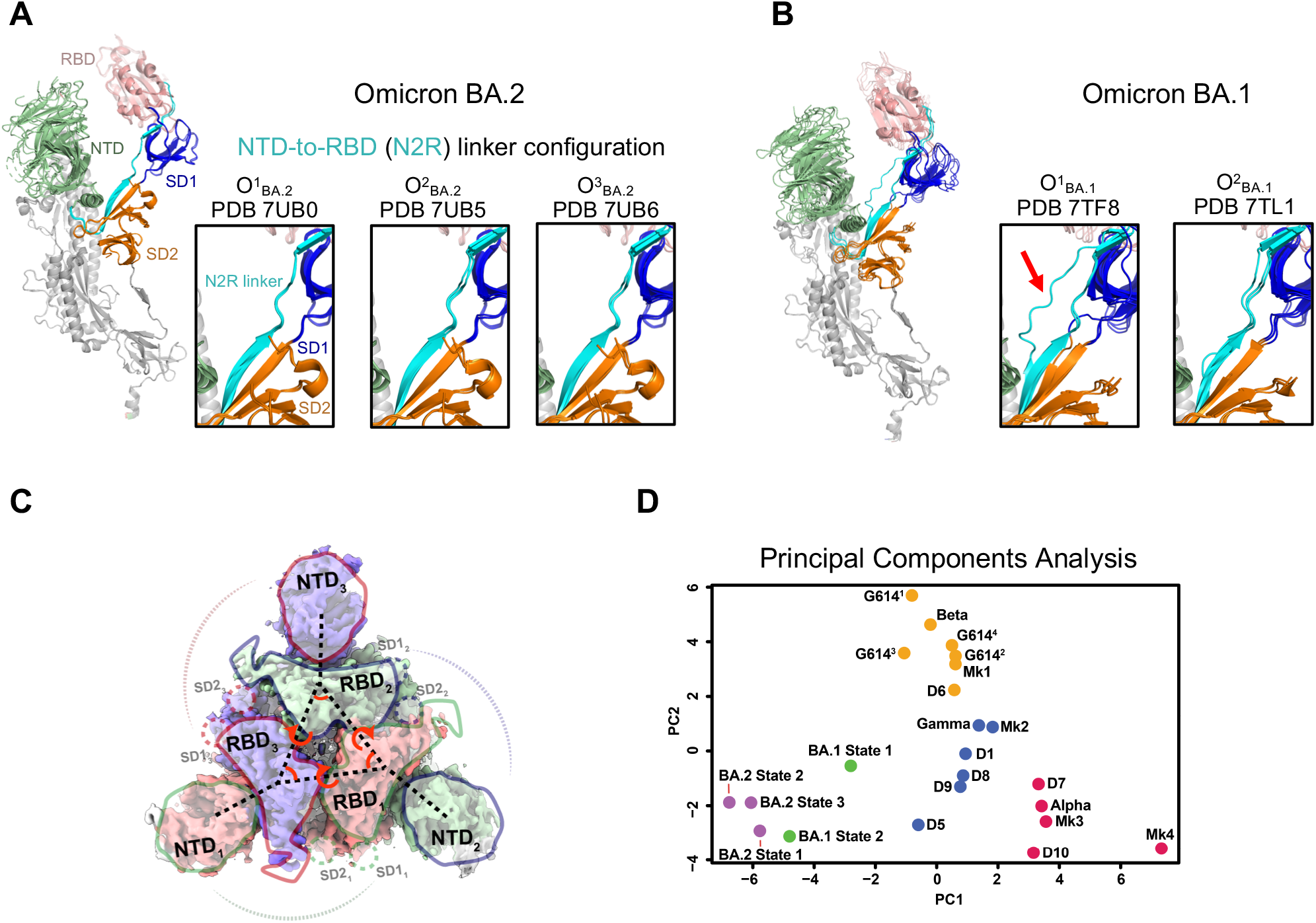
Intra- and Interprotomer communication in the Omicron BA.2 3-RBD-down states, Related to Figure S6. **A**. Omicron BA.2 3-RBD-down structures shown with the three protomers aligned using S2 subunit residues 908-1035. The structures are colored by domain; green: NTD, salmon: RBD, cyan: N2R linker; blue: SD1 subdomain, orange: SD2 subdomain, white: S2 subunit. The insets show zoomed-in views of the N2R region that connects the NTD and RBD within a protomer for the Omicron BA.2 S 3-RBD-down structures O^1^_BA.2_ (PDB: 7UB0), O^2^_BA.2_ (PDB: 7UB5) and O^3^_BA.2_ (PDB: 7UB6). **B**. Same as **A**. but for the Omicron BA.1 3-RBD-down structures O^1^_BA.1_ (PDB: 7TF8) and O^2^_BA.1_ (PDB: 7TL1). The red arrow in panel **B**. is pointing to the N2R rearranged state that was observed in one of the protomers in the O^1^_BA.2_ (PDB 7TF8) structure. **C**. Inter-protomer vectors describing the relationship between the NTDs, RBDs, and subdomains across protomers overlaid on the O^1^_BA.2_ (EMDB: 26433) cryo-EM reconstruction. **D**. Principle components analysis of the interprotomer vector network distances, angle, and dihedrals for SARS-CoV-2 variant structures. Green, yellow, blue, and red points are K means cluster assignments (K=4) for PCA of the dataset excluding the BA.2 variant. Each point represents a variant structure. The variants shown include D614G (G614^1^, G614^2^, G614^3^ and G614^4^), Alpha, Beta, Gamma, a mink-associated variant (Mk1, Mk2, Mk3 and Mk4), BA.1 State 1 (O^1^_BA.1_) and State 2 (O^2^_BA.1_), and BA.2 (O^1^_BA.2_, O^2^_BA.2_, and O^3^_BA.2_).

We previously defined a set of vectors that report on the overall domain organization of the S protein (**Figure 4C**) (Gobeil et al., 2022, Gobeil et al., 2021b, Henderson et al., 2020). Analyzing the BA.2 S 3-RBD-down structures using principal components analysis (PCA) of our previously described interprotomer vectors, we showed that the Omicron BA.2 S 3-RBD-down structures clustered close to the BA.1 S 3-RBD-down structures and were separated from the other variants, including the Alpha, Gamma, Beta, D614G and a mink-associated variant (**Figure 4D**). The three BA.2 3-RBD-down spike structures cluster closely together in a region of the PCA space closest to the O^2^_BA.1_ structure. This is consistent with the observed RBD-RBD packing in both the BA.1 and BA.2 S ectodomain structures.

Taken together our structural studies show that the acquired Omicron BA.2 S mutations lead to further stabilization of the 3-RBD-down state, compared to the BA.1 S protein, through restructuring of the RBD-RBD interface resulting in additional stabilizing interprotomer interactions.

### Antigenicity of the Omicron BA.2 S protein

To assess the antigenic impact of the Omicron BA.2 spike mutations, we tested binding of spike-directed antibodies to two different S protein fragments: a monomeric RBD-only construct and the S protein ectodomain (S-GSAS platform) that were used in our cryo-EM structural analysis, here and in previously published studies (Gobeil et al., 2021a, Gobeil et al., 2021b, Gobeil et al., 2022) (**Figures 5, S11 and S12**). We tested binding of two representative NTD-directed antibodies: the neutralizing antibody DH1050.1 that targets an antigenic supersite in the NTD, and antibody DH1052 that recognizes a different NTD epitope, is non-neutralizing in *in vitro* assays but protects against SARS-CoV-2 challenge in animal models (**Figure S13**) (McCallum et al., 2021, Li et al., 2021). Both NTD-directed antibodies lost binding to the BA.1 and BA.2 S ectodomains. For testing RBD-directed antibodies, we chose representative examples of Receptor Binding Motif (RBM) targeting antibodies (DH1041 and DH1042), RBD inner face targeting antibodies (DH1047, S2X259 and CR3022) and RBD outer face targeting antibodies (DH1044, DH1193 and S309) (**Figures 5A**) (Li et al., 2021, Gobeil et al., 2022, Tortorici et al., 2021, Pinto et al., 2020). The RBM targeting antibodies DH1041 and DH1042 bound the D614G spike but lost binding to both Omicron BA.1 and BA.2 spikes, consistent with the accumulation of escape mutations around the ACE2 binding ridge (**Figure 5B**).

**Figure 5.**
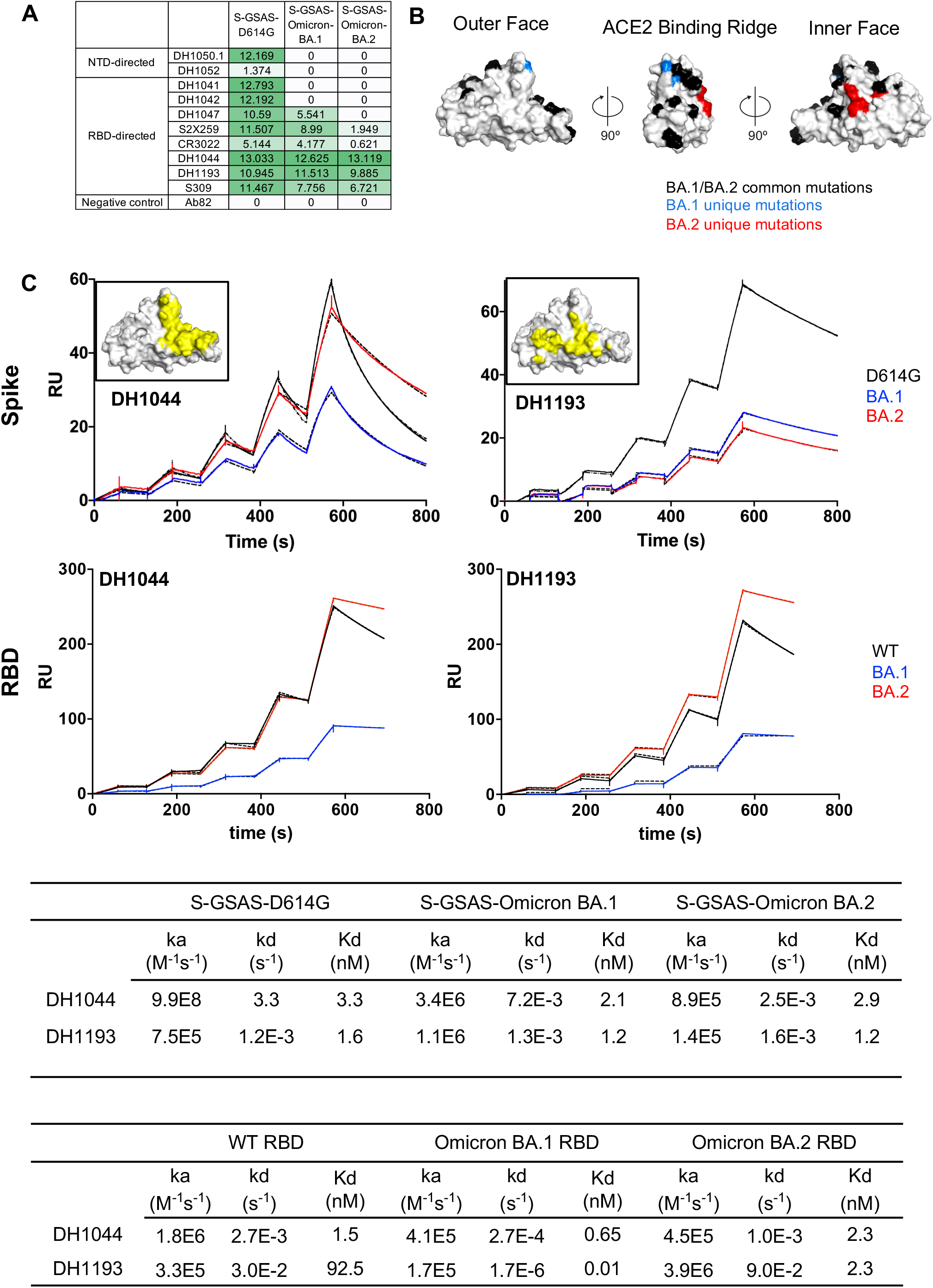
Antigenicity of the SARS-CoV-2 Omicron BA.2 S protein, Related to Figures S11 and S12, and Table S1. **A**. Antibody binding to SARS-CoV-2 S proteins measured by ELISA. The binding values were obtained by calculated area under curve of ELISA binding curves, and are color coded with a dark green to white gradient where dark green indicates tighter binding and white indicates no binding. **B**. Locations of Omicron BA.1 and BA.2 mutations mapped on the RBD surface. **C**. First row, binding of DH1044 and DH1193 Fabs to D614G (black), BA.1 (blue) and BA.2 (red) S protein ectodomains measured by SPR. Second row, binding of antibodies DH1044 and DH1193 to WT (black), Omicron-BA.1 (blue), and Omicron-BA.2 (red) RBD, measured by SPR using single-cycle kinetics. The solid lines are the binding sensorgrams; the dotted lines show fits of the data to a 1:1 Langmuir binding model. Affinity and kinetics of DH1044 and DH1193 Fab binding to the (top) S protein ectodomain, and (bottom) monomeric RBD-only constructs, are tabulated below.

Of the RBD outer face binding antibodies tested, S309 lost ∼40% binding to the BA.2 spike relative to the D614G construct, consistent with the loss of its neutralization efficacy against BA.2 (Iketani et al., 2022, Takashita et al., 2022), although a recent study has reported that although S309 has lost neutralization activity, it still retains protective efficacy against three SARS-CoV-2 Omicron sub-lineages (BA.1, BA.1.1, and BA.2) (Case et al., 2022). We had previously reported two antibodies, DH1044 and DH1193, that bind the outer RBD face and retain neutralizing activity against Omicron BA.1 (Li et al., 2021, Gobeil et al., 2022). Both antibodies retained nM binding affinity to the BA.2 RBD-only and S protein ectodomain constructs (**Figures 5C**) and effectively neutralized Omicron BA.1, BA.2 and BA.3 in a pseudovirus neutralization assay (**Table S1**). We observed that DH1044 binding to the D614G, BA.1 and BA.2 spikes followed the same trend as DH1044 binding to the corresponding RBD-only constructs. In contrast, DH1193 bound the BA.2 RBD-only construct at higher levels than it did the WT RBD construct, while the BA.2 S ectodomain bound DH1193 at a substantially lower relative to the D614G spike (**Figure 5C**). This is likely due to conformational effects that occur in the context of the spike and are not present in the RBD-only context. As DH1193 binds to an RBD up conformation, its binding level to the BA.2 spike may be diminished due to the lower propensity of the BA.2 spike to adopt the RBD up configuration. DH1044, on the other hand, binds the RBD-down state and, thus, its epitope is not similarly constrained by spike conformational dynamics.

One of the most dramatic antigenic consequences of the Omicron BA.2 mutations is the elimination of Class 4 antibody neutralizing activity (Barnes et al., 2020, Iketani et al., 2022). Of note, antibody S2X259 retains activity against Omicron BA.1 but is unable to neutralize BA.2 (Iketani et al., 2022, Cameroni et al., 2021, Tortorici et al., 2021). Examining the RBD binding interface of S2X259 (**Figure 6**), we found that the restructuring of the 371-376 RBD loop in the BA.2 S protein resulted in a clash with the bound antibody. This was reflected in the binding of S2X259 to the Omicron S proteins (**Figure 5A**), where substantial binding was retained with the BA.1 S protein but not to BA.2 S. Similar binding trends were also observed with the other Class 4 antibodies CR3022 and DH1047 (**Figure 6**) (Li et al., 2021, Yuan et al., 2020, Martinez et al., 2021). Like S2X259, the epitopes of both DH1047 and CR3022 was disrupted by the restructuring of the 371-376 RBD loop.

**Figure 6.**
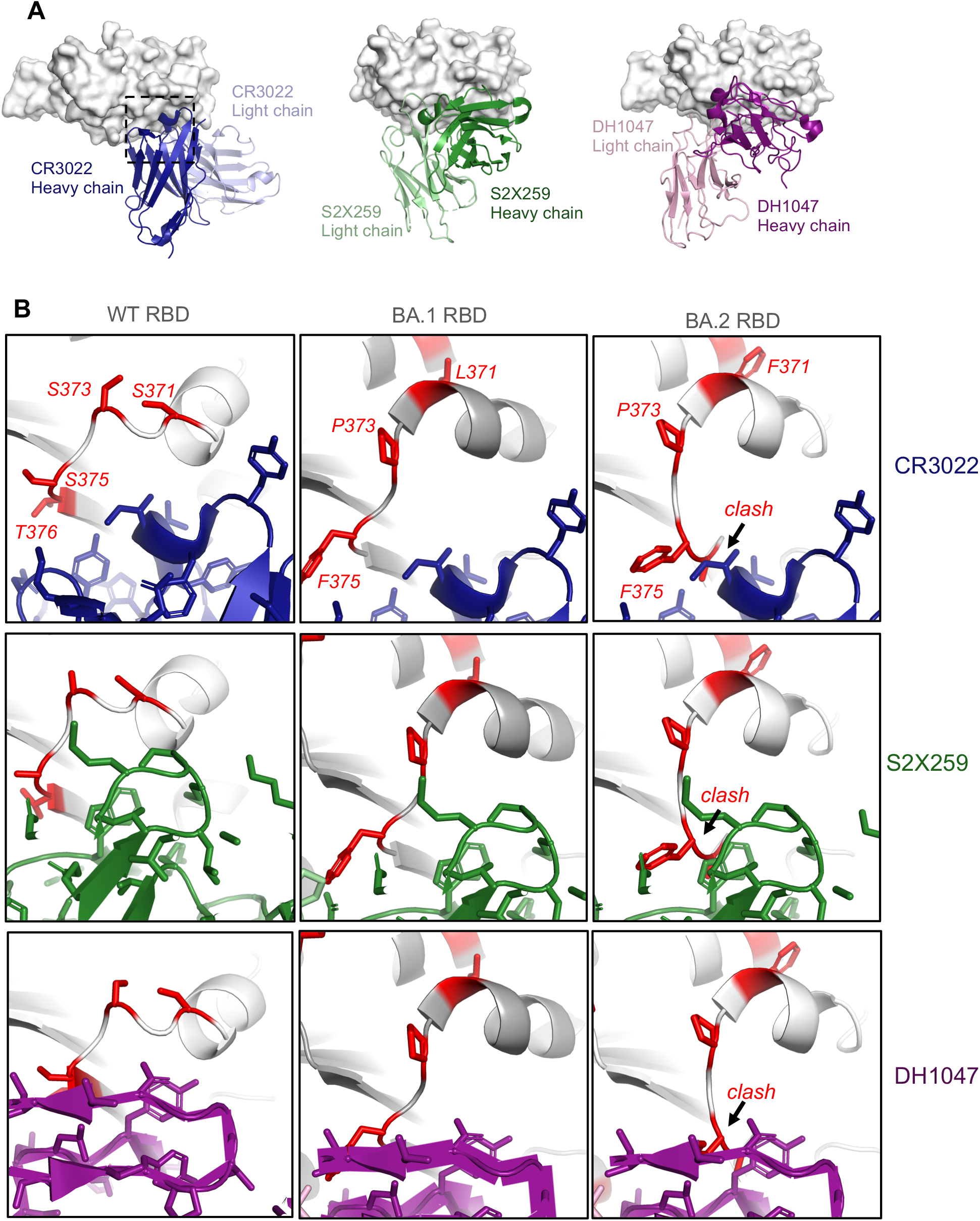
Structural basis for loss in binding of Class 4 RBD binding antibodies to Omicron BA.2 S protein. Related to Figures S11 and S12. **A**. Left. Crystal structure of CR3022 bound to SARS-CoV-2 WT RBD (PDB: 7LOP), with the RBD shown as grey surface, and CR3022 heavy and light chains colored dark blue and light blue, respectively. The dotted square indicates the zoomed-in areas shown in **B**. Middle. Cryo-EM structure of of S2X259 bound to SARS-CoV-2 WT RBD (PDB: 7RAL), with the RBD shown as grey surface, and S2X259 heavy and light chains colored dark green and light green, respectively. Right. Cryo-EM structure of of DH1047 bound to SARS-CoV-2 WT RBD (PDB: 7LD1), with the RBD shown as grey surface, and DH1047 heavy and light chains colored dark magenta and light pink, respectively. **B**. Zoomed in images of antibodies CR3022 (blue; PDB: 7LOP), S2X259 (green; PDB: 7RAL), and DH1047 (magenta; PDB: 7LD1), bound to WT RBD (leftmost panels). The middle and right panels show models of the antibodies bound to BA.1 (PDB: 7TF8) and BA.2 (PDB: 7UBO) RBDs. The models were prepared by aligning the variant RBDs with the antibody-bound RBD in each structure. For the WT RBD, residues that are mutated in the Omicron BA.2 variant are colored red and shown as sticks. For the BA.1 and BA.2 RBDs, the mutated residues in each are colored red. A region where the RBD 371-376 loop clashes with the antibody is indicated in the BA.2 RBD-bound models.

In summary, these results show that the acquired mutations in the Omicron BA.2 S protein affect the binding of RBD-directed antibodies by conformational effects related to the RBD up/down transitions in the context of the spike as well as by conformational changes within the RBD itself.

### Conformational changes in the S2 subunit of the Omicron BA.2 S protein

Omicron BA.1 and BA.2 S proteins share all but two S2 subunit mutations; the N865K and L981F substitutions in BA.1 do not occur in the BA.2 spike (**Figures 1A and 7A**). Within the S2 subunit is a quaternary glycan cluster that binds to Fab-dimerized, glycan-reactive (FDG) antibodies (Williams et al., 2021). We found similar levels of binding of FDG antibodies 2G12 and DH851.3 to the Omicron BA.1 and BA.2 S proteins (**Figure 6B**). Given the previously observed sensitivity of FDG antibody binding to S2 subunit conformational changes (Edwards et al., 2021, Gobeil et al., 2022), this suggested that the differences at S2 residue positions 856 and 981 do not cause substantial changes to the overall conformation of the pre-fusion BA.2 spike S2 subunit.

**Figure 7.**
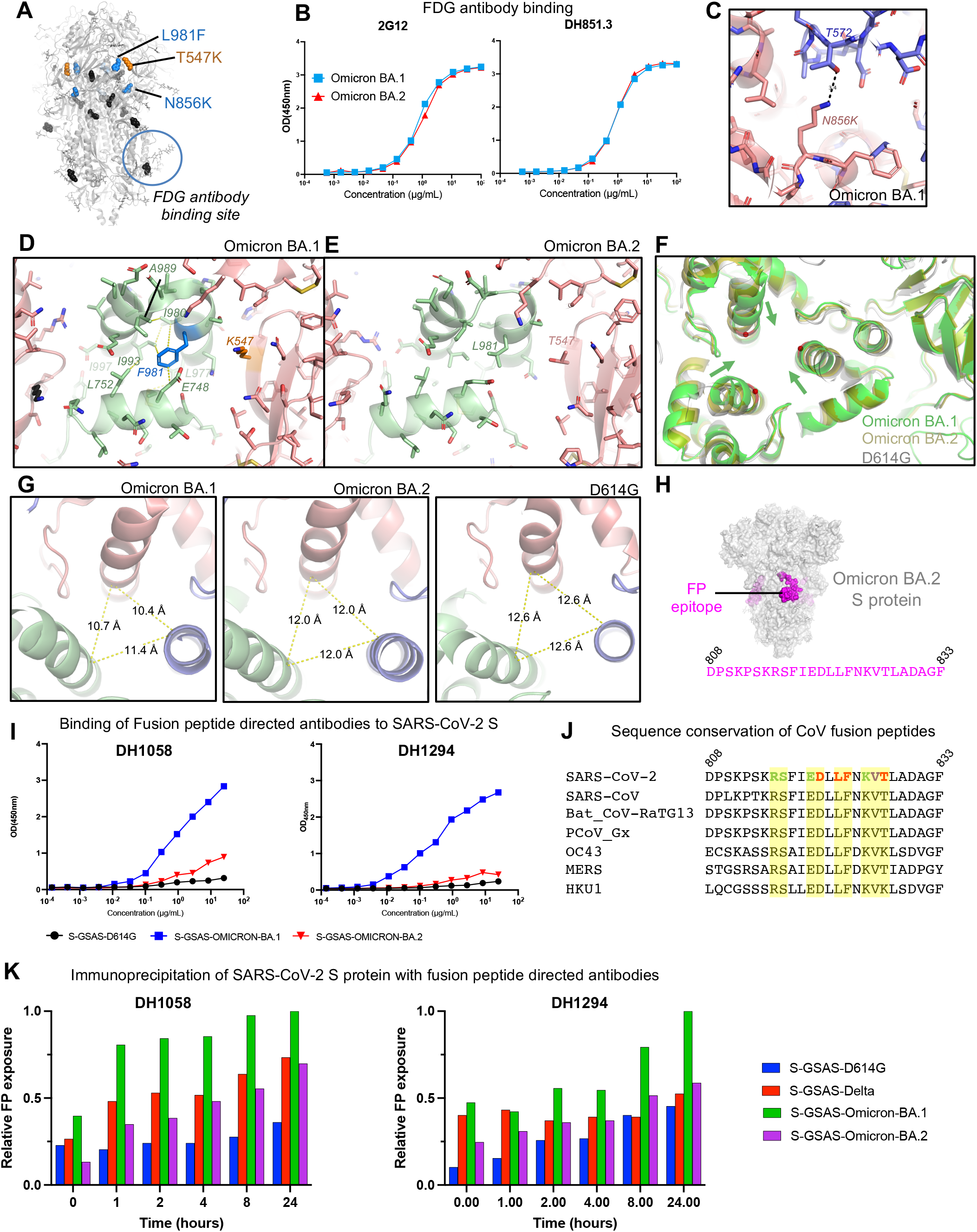
Conformation and antigenicity of the S2 subunit of the Omicron BA.2 S protein, Related to Figures S14 and S15. **A**. Omicron BA.1 spike shown in grey with S2 and SD1 mutations shown as spheres. The S2 mutations that BA.1 and BA.2 have in common are colored black. The L981F and N856K substitutions that occur in the BA.1 but not in the BA.2 spike are colored blue, and the SD1 T547K substitution that occurs in BA.1 but not in the BA.2 spike is colored orange. The glycan cluster that binds Fab-dimerized glycan-reactive (FDG) antibodies is marked with a circle. **B**. Binding of FDG antibodies 2G12 and DH851.3 to Omicron BA.1 and BA.2 S proteins. **C**. Zoom-in of the region around the N856K substitution in the BA.1 spike showing an interprotomer hydrogen bond between K856 and T572. **D**. Zoom-in of the region around the L981F and T547K substitutions in the Omicron BA.1 spike. The yellow dotted lines indicate van der Waals contacts. **E**. Same as is D but for the Omicron BA.2 spike. **F**. Overlay of the D614G, BA.1 and BA.2 spikes showing the movement of the BA.1 S2 subunit helices towards the center of the trimer axis. **G**. View of the S2 subunit helices, showing interprotomer distances between the Ca atom of residue R995, which was used to measure the differences in the arrangement of this region between the different spikes. **H**. Omicron BA.2 spike with the location of the fusion peptide shown in magenta. The sequence below spans the magenta regions mapped on the structure. **I**. ELISA binding of antibodies (left) DH1058 and (right) DH1294 to the D614G (black), BA.1 (blue) and BA.2 (red) S protein ectodomains **J**. Sequence alignment of the fusion peptide region in diverse CoV S proteins. In the SARS-CoV-2 sequence, the colored residues indicate contacts observed with FP-directed antibody DH1058 in the crystal structure of FP bound to DH1058 (PDB: 7TOW). Red indicates residues that make both main chain and side chain contacts, green indicates residues that only make side chain contacts, and purple indicates residues that contact DH1058 only through the main chain. Contacts indicated here include both direct contact with the antibody as well as water-mediated contacts. **K**. Time dependent exposure of fusion peptide (FP) to FP-directed antibodies, DH1058 and DH1294. Shown as relative FP exposure ranging from 0 to 24 hours.

We next examined the local regions around the mutations that were different between BA.1 and BA.2. The N856K substitution in BA.1 introduced an interprotomer hydrogen bond involving the side chains of K856 and T572 (**Figure 7C**). The absence of this mutation in BA.2 could lead to local destabilization in this region relative to BA.1. The L981F mutation in the BA.1 S protein occurs in a structurally important region, proximal to residues K986 and V987 at the junction of the Heptad Repeat 1 (HR1) and Central Helix (CH), where engineering two consecutive prolines, the commonly used “2P” mutations, blocks the transition from pre- to post-fusion conformation (Wrapp et al., 2020, Pallesen et al., 2017). In the Omicron BA.1 structure, F981 inserts into a pocket between three interprotomer helices, and in doing so, sets up an extensive van der Waals interaction network involving residues E748, L752, L977, I980, A989, I993, and I997 (**Figure 7D**). Interestingly, the lone T547K SD1 subdomain mutation in BA.1 (not present in BA.2) is next to the helical segment that contains residue F981, with the K547 side chain stacked against this helix. The less bulky L981 in the Omicron BA.2 spike cannot mediate the van der Waals network that F981 does in the BA.1 S protein (**Figure 7E**). We asked if there were any changes in the local region around this mutation, and found that the CH helices in the Omicron BA.1 spike are pulled towards the central trimer axis relative to the positions of these helixes in the D614G and BA.2 S proteins (**Figure 7F**). To quantify the extent of this shift, we measured the distances between Cα of residue 995 (**Figure 7G**) and found that the D614G spike helices showed the greatest separation with the 995 Cαs 12.6 Å apart. The Omicron BA.2 helices are slightly closer together with a separation of 12 Å between the residue 995 Cα atoms, whereas the Omicron BA.1 helices were on average ∼2 Å closer, with the distances showing more asymmetry with values of 10.4 Å, 10.7 Å, and 11.4 Å. As noted, these local changes did not percolate globally through the structure as conformation sensitive recognition of the glycan patch by FDG antibodies remained unchanged (**Figures 7A and 7B**).

We next probed the conformation of the fusion peptide (FP) region using FP-directed antibodies, DH1058 and DH1294 (Li et al., 2021, Gobeil et al., 2021b, Gobeil et al., 2022) (**Figures 7H-K** and **S14-S15**). Both antibodies mapped by ELISA to a 25 amino acid peptide spanning the SARS-CoV-2 FP region (**Figures 7H and S14**) (Li et al., 2021, Gobeil et al., 2021b). Like DH1058, DH1294 bound diverse CoV spikes (**Figure S14**). Analysis of the FP sequences in these diverse spikes revealed striking conservation of the residues that were identified as DH1058 antibody contacts in a crystal structure of DH1058 bound to the FP (PDB 7TOW) (**Figure 7J**) (Gobeil et al., 2022). We had previously observed increased accessibility of the Omicron BA.1 FP (relative to the FP in D614G and Delta variant spikes) to DH1058 binding by ELISA (Gobeil et al., 2022). We observed similar enhanced binding of the Omicron BA.1 spike to DH1294, while both DH1058 and DH1294 bound the Omicron BA.2 spike at much lower levels (**Figure 7I**), suggesting that the BA.2 FP was less accessible to the FP-directed antibodies compared to the BA.1 FP. Immunoprecipitation assays showed time-dependent enhancement of binding upon incubation of the FP-directed antibodies with SARS-CoV-2 S ectodomain constructs, with Omicron BA.1 showing the highest levels of binding (**Figures 7K** and **S15**).

In summary, our structural analysis and binding data reveal differences in the S2 subunit between the Omicron BA.1 and BA.2 S proteins, including differences in the configuration of the critical HR1-CH region and altered fusion peptide accessibility.

## Discussion

The emergence of the SARS-CoV-2 Omicron variant triggered widespread alarm due to its highly mutated spike protein (Viana et al., 2022). The rapid spread of the Omicron variant was at first dominated by its BA.1 sub-lineage (**Figure S1**). In time, a second Omicron sub-lineage, BA.2, started spreading and has now overtaken the BA.1 in several locations. As the spike (S) protein is central to several properties that influence the spread of a variant, here we determined structures of the BA.2 spike to compare them to our previously determined structures of the Omicron BA.1 S protein, aiming to arrive at a structure-guided understanding of the similarities and differences between the S protein-driven properties of these two phylogenetically related sub-lineages.

A unifying structural feature of the Omicron variant is the tight packing of its RBDs in the 3-RBD-down (or closed) spike where the receptor binding site and many immunodominant epitopes are inaccessible. We previously recognized that an interfacial RBD loop in the BA.1 S protein containing three mutations – S371L, S373P and S375F – mediates interprotomer RBD-RBD packing in the closed Omicron BA.1 S protein (Gobeil et al., 2022). The BA.2 S protein incorporates additional strategically positioned mutations in this interfacial RBD loop leading to restructuring of this region and facilitating both improved internal packing of the RBD core as well as improved packing of the RBD-RBD interface in the 3-RBD-down spike. We expect the spike proteins of the BA.3, BA.4 and BA.5 Omicron variant sub-lineages will also have similarly tightly packed 3-RBD-down states, as they all contain the key S371F, S373P and S375F mutations of BA.2. In addition, the BA.4 and BA.5 spikes also include the T376A substitution that occurs in the BA.2 spike (but not in BA.1 and BA.3). The increased stabilization of the closed state may result in low immunogenicity of the spike, which may translate to less effective protection from an Omicron infection for unvaccinated individuals without prior infection by any other variant. For the same reason a vaccine based on the Omicron spike may exhibit lowered immunogenicity as large proportions of the immunogenic RBD sites would be occluded. This mutation-induced restructuring of intra- and inter-RBD structures contributes to the higher immune evasion observed in BA.2, particularly the loss in activity of Class 4 RBD-directed antibodies (Barnes et al., 2020, Iketani et al., 2022), where we showed that a combination of local structural effects caused by the RBD mutations, as well as spike conformational effects may play a role in the dramatic loss in activity observed.

Even though the BA.2 spike had a better packed RBD-RBD interaction in the closed state than the BA.1 spike, they both bound similar levels of ACE2 receptor in our *in vitro* ELISA assay suggesting that the receptor binding competencies remain similar between the two variants despite their modified interprotomer associations. We observed differences between the BA.1 and BA.2 spikes in accessibility of the fusion peptide (FP). Our results demonstrating that these FP-directed antibodies can induce FP exposure provides intriguing insights into the mechanism of action of these broadly reactive antibodies and the dynamics of this functionally critical and highly conserved S2 subunit region. While FP-directed antibodies like DH1058 and DH1294 are non-neutralizing, it remains to be determined whether they are protective *in vivo*. Being able to effectively target these highly conserved regions could lead to strategies to elicit broad protective responses to counteract future variants or CoV outbreaks.

The key differences in the BA.1 and BA.2 spikes include (1) better reinforced RBD-RBD-packing in the BA.2 closed state, (2) better internal packing of the BA.2 RBD, and (3) a less accessible fusion peptide. All these factors, as well as the absence of the N2R rearranged state observed in the BA.1 3-RBD-down O^1^_BA.1_ structure, suggest a more stable architecture of the BA.2 spike compared to the BA.1 spike. Over the course of SARS-CoV-2 evolution, particularly in variants that have emerged to become VOCs, we have observed the S proteins balance between stability of the pre-fusion state with features such as greater propensity to go to a more open state that may lead to greater transmissibility (Gobeil et al., 2021b, Zhang et al., 2021, Zhang, 2022, Zhou, 2021). In the BA.2 spike we see the balance tilt towards greater stability relative to BA.1. It is tempting to speculate that the structural features that make the S protein in BA.1 less stable than in BA.2, including increased propensity for the RBD to transition to the “up” state and more facile release of the fusion peptide, may have contributed to its rapid spread during the start of the Omicron wave, while the greater stability of the BA.2 spike and its increased immune evasion properties may be allowing it to make its current gains in the wake of a receding BA.1 wave.

Overall, our studies uncover defining characteristics of the Omicron variant and through atomic level structural determination assign purpose to key mutations. They further highlight striking differences in functional regions between the BA.1 and BA.2 sub-lineages that could be responsible for differences in their biology.

## Acknowledgements

Cryo-EM data were collected at the Duke Krios at the Duke University Shared Materials Instrumentation Facility (SMIF), a member of the North Carolina Research Triangle Nanotechnology Network (RTNN), which is supported by the National Science Foundation (award number ECCS-2025064) as part of the National Nanotechnology Coordinated Infrastructure (NNCI). This study utilized the computational resources offered by Duke Research Computing (http://rc.duke.edu; NIH 1S10OD018164-01) at Duke University. This work was supported by NIH R01 AI145687 (P.A. and W.W.), and AI165147 (P.A. and R.H.), NIH, NIAID, DMID grant P01 AI158571 (B.F.H.).

## Author contributions

V.S., and P.A. determined cryo-EM structures, and prepared the first draft of the manuscript. J.J.L. purified RBD and spike proteins, and performed SPR experiments. V.S., S.M-C.G., R.C.H, A.M., and P.A. analyzed structures. S.M-C.G. performed immunoprecipitation assays. R.P., M.B., M.D., and M.M. performed ELISA assays. K.J., X.H., A.M., M.S. and E.B. expressed and purified proteins. R.J.E. performed NSEM analysis. K.W., and W.W. analyzed data. K.O.S., B.C., and X.L. provided key reagents. B.F.H provided S RBD and S2 antibodies, supervised ELISA assays, and edited the manuscript. All authors reviewed and approved the manuscript. P.A. supervised the study and reviewed all data.

## STAR METHODS

### RESOURCE AVAILABILITY

#### Lead Contact

Further information and requests for resources and reagents should be directed to and will be fulfilled by the Lead Contact, Priyamvada Acharya.

#### Materials Availability

Further information and requests for resources and reagents should be directed to Priyamvada Acharya. Plasmids generated in this study have been deposited to Addgene with the following accession numbers: 184829, 184830, 184831.

#### Data and Code Availability

- Cryo-EM reconstructions and atomic models generated during this study are available at wwPDB and EMBD (https://www.rcsb.org; http://emsearch.rutgers.edu) under the accession codes PDB IDs 7UB0, 7UB5, 7UB6, and EMDB IDs 26433, 26435, 26436, 26643 and 26647.
- This paper does not report original code.
- Any additional information required to reanalyze the data reported in this paper is available from the lead contact upon request.

### EXPERIMENTAL MODEL AND SUBJECT DETAILS

#### Cell culture

Gibco FreeStyle 293-F cells (embryonal, human kidney) were incubated at 37°C and 9% CO_2_ in a humidified atmosphere. Cells were incubated in FreeStyle 293 Expression Medium (Gibco) with agitation at 120 rpm. Plasmids were transiently transfected into cells using Turbo293 (SpeedBiosystems) and incubated at 37 °C, 9% CO2, 120 rpm for 6 days. On the day following transfection, HyClone CDM4HEK293 media (Cytiva, MA) was added to the cells. Antibodies were produced in Expi293 cells (embryonal, human kidney). Cells were incubated in Expi293 Expression Medium at 37°C, 120 rpm and 8% CO_2_ in a humidified atmosphere. Plasmids were transiently transfected into cells using the ExpiFectamine 293 Transfection Kit and protocol (Gibco).

### METHOD DETAILS

#### Plasmids

Site-directed mutagenesis were performed and sequences confirmed by GeneImmune Biotechnology (Rockville, MD). The SARS-CoV-2 spike protein ectodomain constructs comprised the S protein residues 1 to 1208 (GenBank: MN908947) with the D614G mutation, the furin cleavage site (RRAR; residue 682-685) mutated to GSAS, a C-terminal T4 fibritin trimerization motif, a C-terminal HRV3C protease cleavage site, a TwinStrepTag and an 8XHisTag. All spike ectodomains were cloned into the mammalian expression vector pαH. The WT RBD construct obtained from BEI resources (Catalog number NR-52309) expressed the receptor binding domain (RBD) of the spike (S) glycoprotein gene from severe acute respiratory syndrome-related coronavirus 2 (SARS-CoV-2), Wuhan-Hu-1 (GenBank: MN908947) with an N-terminal S protein signal sequence to the spike RBD (amino acids 319 to 541) and a C-terminal 6X-histidine tag. The sequence was codon optimized for mammalian expression and subcloned into the pCAGGS mammalian expression vector under the AG promoter. All plasmids generated in this study have been deposited to Addgene (https://www.addgene.org).

#### Protein purification

Spike ectodomains were harvested from the concentrated supernatant on day 6 post transfection. The spike ectodomains were purified using StrepTactin resin (IBA LifeSciences) and size exclusion chromatography (SEC) using a Superose 6 10/300 GL Increase column (Cytiva, MA) equilibrated in 2mM Tris, pH 8.0, 200 mM NaCl, 0.02% NaN_3_. All purification steps were performed at room temperature within a single day. Protein quality was assessed by SDS-PAGE using NuPage 4-12% (Invitrogen, CA). The purified proteins were flash frozen and stored at -80°C in single-use aliquots. Each aliquot was thawed by a 20-minute incubation at 37 °C before use.

RBD variants were harvested from concentrated supernatant on the 6th day post transfection. The RBDs containing an 6x-Histidine tag were purified via nickel affinity chromatography using a HisTrap excel column (Cytiva, MA). Concentrated supernatant was loaded onto the column and the column washed with buffer A (1x PBS pH 8.0) until baseline. The proteins were eluted from the column by applying a gradient over 40 CV from 100% buffer A to 100% buffer B (1x PBS pH 8.0, 1 M Imidazole). Fractions containing RBD were pooled, concentrated, and further purified by size exclusion chromatography (SEC) using a Superdex 200 Increase 10/300 GL column (Cytiva, MA) equilibrated with 1x PBS, pH 8.0. All steps of the purification were performed at room temperature. Protein quality was assessed by SDS-PAGE using NuPage 4-12% (Invitrogen, CA). The purified proteins were flash frozen and stored at - 80°C in single-use aliquots. Each aliquot was thawed at 4 °C before use.

Antibodies were produced in Expi293F cells and purified by Protein A affinity and digested using LysC to generate Fab fragments. ACE2 with human Fc tag was purified by Protein A affinity chromatography and SEC.

#### Differential scanning fluorimetry

DSF assays were performed using Tycho NT.6 (NanoTemper Technologies). Spike ectodomains were diluted to approximatively 0.15 mg/ml. Intrinsic fluorescence was measured at 330 nm and 350 nm while the sample was heated from 35 to 95 °C at a rate of 30°C/min. The ratio of fluorescence (350/330 nm) and inflection temperatures (T_i_) were calculated using the inbuilt software in the Tycho NT. 6.

#### ELISA assays

Spike ectodomains were tested for antibody- or ACE2-binding in ELISA assays as previously described (Edwards et al., 2021). Serially diluted spike protein was bound in wells of a 384-well plates, which were previously coated with streptavidin (Thermo Fisher Scientific, MA) at 2 µg/mL and blocked. Proteins were incubated at room temperature for 1 hour, washed, then human mAbs were added at 10 µg/ml. Antibodies were incubated at room temperature for 1 hour, washed and binding detected with goat anti-human-HRP (Jackson ImmunoResearch Laboratories, PA) and TMB substrate.

Recombinant FDG mAbs were tested for binding to the SARS-CoV-2 spikes in ELISA. Briefly, spike proteins (20ng) were captured by streptavidin (30ng per well) to individual wells of a 384-well Nunc-absorb ELISA plates using PBS-based buffers and assay conditions as previously described (PMID: 34019795; PMID: 28298421; PMID: 28298420). Commercially obtained D-mannose (Sigma, St. Louis, MO) was used to outcompete mAb binding to glycans on the spike proteins; D-mannose solutions were also produced in ELISA PBS-based glycan buffers at a concentration of [1M] D-mannose as described (PMID: 34019795). Mouse anti-monkey IgG-HRP (Southern Biotech, CAT# 4700-05) and Goat anti-human IgG-HRP (Jackson ImmunoResearch Laboratories, CAT# 109-035-098) secondary antibodies were used to detect antibody bound to the spike proteins. HRP detection was subsequently quantified with 3,30,5,50-tetramethylbenzidine (TMB) by measuring binding levels at an absorbance of 450nm, and binding titers were also reported as Log area under the curve (AUC).

#### Surface Plasmon Resonance

Binding experiments were performed using SPR on a Biacore T-200 (Cytiva, MA, formerly GE Healthcare) with HBS buffer supplemented with 3 mM EDTA and 0.05% surfactant P-20 (HBS-EP+, Cytiva, MA). All binding assays were performed at 25 °C.

Spike variants were captured on a Series S Strepavidin (SA) chip (Cytiva, MA) coated at 200 nM (120s at 5µL/min). Fabs were injected at concentrations ranging from 0.5 nM to 8 nM (prepared in a 2-fold serial dilution manner) over the S proteins using the single cycle kinetics mode with 5 concentrations per cycle. The surface was regenerated after the last injection with 3 pulses of a 50mM NaoH + 1M NaCl solution for 10 seconds at 100µL/min.

RBD binding to IgG’s was assessed using a Series S CM5 chip (Cytiva, MA) which was labeled with anti-human IgG (fc) antibody using a Human Antibody Capture Kit (Cytiva, MA). IgGs were then coated at 200 nM (120s at 5 µL/min). RBDs were injected at concentrations ranging from 0.5 nM to 40 nM (prepared in a 2-fold serial dilution manner) over the antibodies using the single cycle kinetics mode with 5 concentrations per cycle. The surface was regenerated after the last injection with 3 pulses of a 3 M MgCl_2_ solution for 10 seconds at 100µL/min. Sensogram data were analyzed using the BiaEvaluation software (Cytiva, MA)

#### Cryo-EM

Purified SARS-CoV-2 spike ectodomains were diluted to a concentration of ∼1.5 mg/mL in 2 mM Tris pH 8.0, 200 mM NaCl and 0.02% NaN_3_ and 0.5% glycerol was added. A 2.3-µL drop of protein was deposited on a Quantifoil-1.2/1.3 grid (Electron Microscopy Sciences, PA) that had been glow discharged for 10 seconds using a PELCO easiGlow™ Glow Discharge Cleaning System. After a 30-second incubation in >95% humidity, excess protein was blotted away for 2.5 seconds before being plunge frozen into liquid ethane using a Leica EM GP2 plunge freezer (Leica Microsystems). Frozen grids were imaged using a Titan Krios (Thermo Fisher) equipped with a K3 detector (Gatan). The cryoSPARC (Punjani et al., 2017) software was used for data processing. Phenix (Liebschner et al., 2019, Afonine et al., 2018), Coot (Emsley et al., 2010), Pymol (Schrodinger, 2015), Chimera (Pettersen et al., 2004), ChimeraX (Goddard et al., 2018) and Isolde (Croll, 2018) were used for model building and refinement.

#### Vector Based Structure Analysis

Vector analysis of intraprotomer domain positions was performed as described previously (Henderson et al., 2020) using the Visual Molecular Dynamics (VMD) (Humphrey et al., 1996) software package Tcl interface. Alpha carbons of each S1 protomer domain and the S2 CD and a S2 sheet motif were used to determine domain centroids for vector calculations. Vectors connecting structurally related domain centroids were used to calculate relevant distances, angles, and dihedrals. Principal components analysis and K-means clustering of the vector sets was performed in R (Team, 2017). Data were centered and scaled for the PCA analyses. Principal components analysis, K-means clustering, and Pearson correlation (confidence interval 0.95, p < 0.05) analysis of vectors sets was performed in R. Data were centered and scaled for the PCA analyses.

#### Difference distance matrices (DDM)

DDM were generated using the Bio3D package (Grant et al., 2020) implemented in R (R Core Team (2014). R: A language and environment for statistical computing. R Foundation for Statistical Computing, Vienna, Austria. URL http://www.R-project.org/)

#### Pseudovirus Neutralization Assay

The pseudovirus neutralization assay performed at Duke has been described in detail (Gilbert et al., 2022) and is a formally validated adaptation of the assay utilized by the Vaccine Research Center; the Duke assay is FDA approved for D614G. For measurements of neutralization, pseudovirus was incubated with 8 serial 5-fold dilutions of antibody samples (1:20 starting dilution using antibodies diluted to 1.0 mg/ml) in duplicate in a total volume of 150 µl for 1 hr at 37°C in 96-well flat-bottom culture plates. 293T/ACE2-MF cells were detached from T75 culture flasks using TrypLE Select Enzyme solution, suspended in growth medium (100,000 cells/ml) and immediately added to all wells (10,000 cells in 100 µL of growth medium per well). One set of 8 wells received cells + virus (virus control) and another set of 8 wells received cells only (background control). After 71-73 hrs of incubation, medium was removed by gentle aspiration and 30 µl of Promega 1X lysis buffer was added to all wells. After a 10-minute incubation at room temperature, 100 µl of Bright-Glo luciferase reagent was added to all wells. After 1-2 minutes, 110 µl of the cell lysate was transferred to a black/white plate. Luminescence was measured using a GloMax Navigator luminometer (Promega). Neutralization titers are the inhibitory dilution (ID) of serum samples at which RLUs were reduced by 50% (ID50) compared to virus control wells after subtraction of background RLUs. Serum samples were heat-inactivated for 30 minutes at 56°C prior to assay.

#### Immunoprecipitation assays

For each time point and controls, 20 µL of 50% slurry Protein A-agarose resin was spun down for 3 minutes at 500 G. The supernatant was then removed and 3 washes with 100µL of PBS were performed with spins of 3 minutes at 3,000G to precipitate the resin. 10µL of PBS was then added to the resin, followed by 5µg of the antibody (DH1058, DH1294, or PGT145 for the control) and 5µg of each spike protein. The combinations were incubated at room temperature according to the time points. To stop the reaction, following 3 washes of the resin with 100µL of PBS as described above, the washed resin was resuspended in 10µL of 4x loading buffer (BioRad) containing DTT and boiled at 95°C for 5 minutes. The samples were then flash-frozen until ready to be loaded on an SDS-PAGE using NuPage 4-12% (Invitrogen, CA). Gel staining was done using the SimplyBlue SafeStain (ThermoFisher) and gel imaging was done using a ChemiDoc (BioRad). Quantification of the band intensity was done using the BioRad Image Lab Software.

### QUANTIFICATION AND STATISTICAL ANALYSIS

<u>No statistical analyses were performed in this study.</u>

