## Supplemental Data for "Cryo-EM structures of SARS-CoV-2 Omicron BA.2 spike"

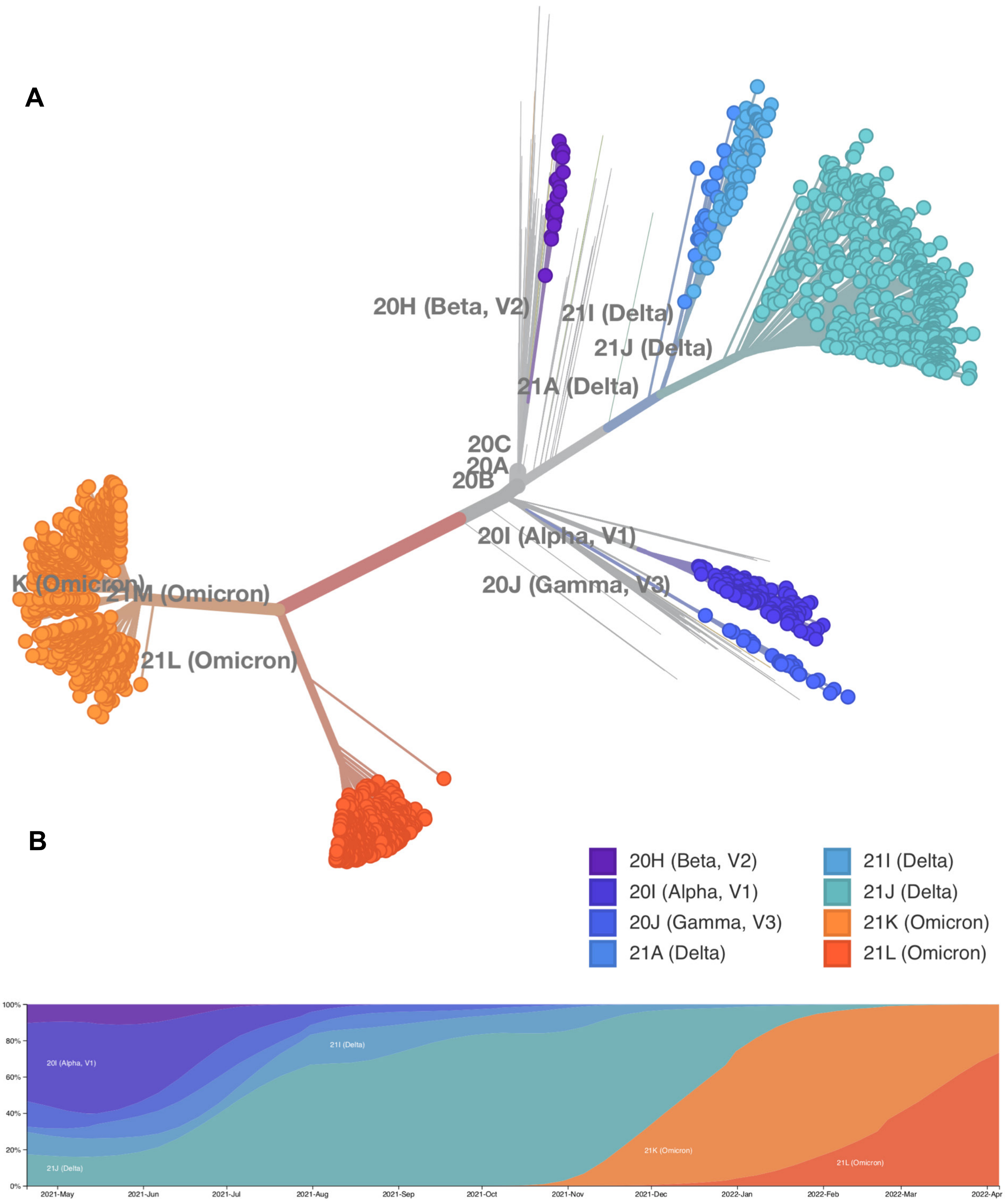


**Fig. S1. Emergence and spread of the SARS-CoV-2 Omicron variant lineages. Related to Figure 1. (A)** Phylogenetic tree of SARS-CoV-2 variants of concern. **(B)** Global frequencies of SARS-CoV-2 variants plotted against time from April 2021 to April 2022 (updated 04/14/2022).

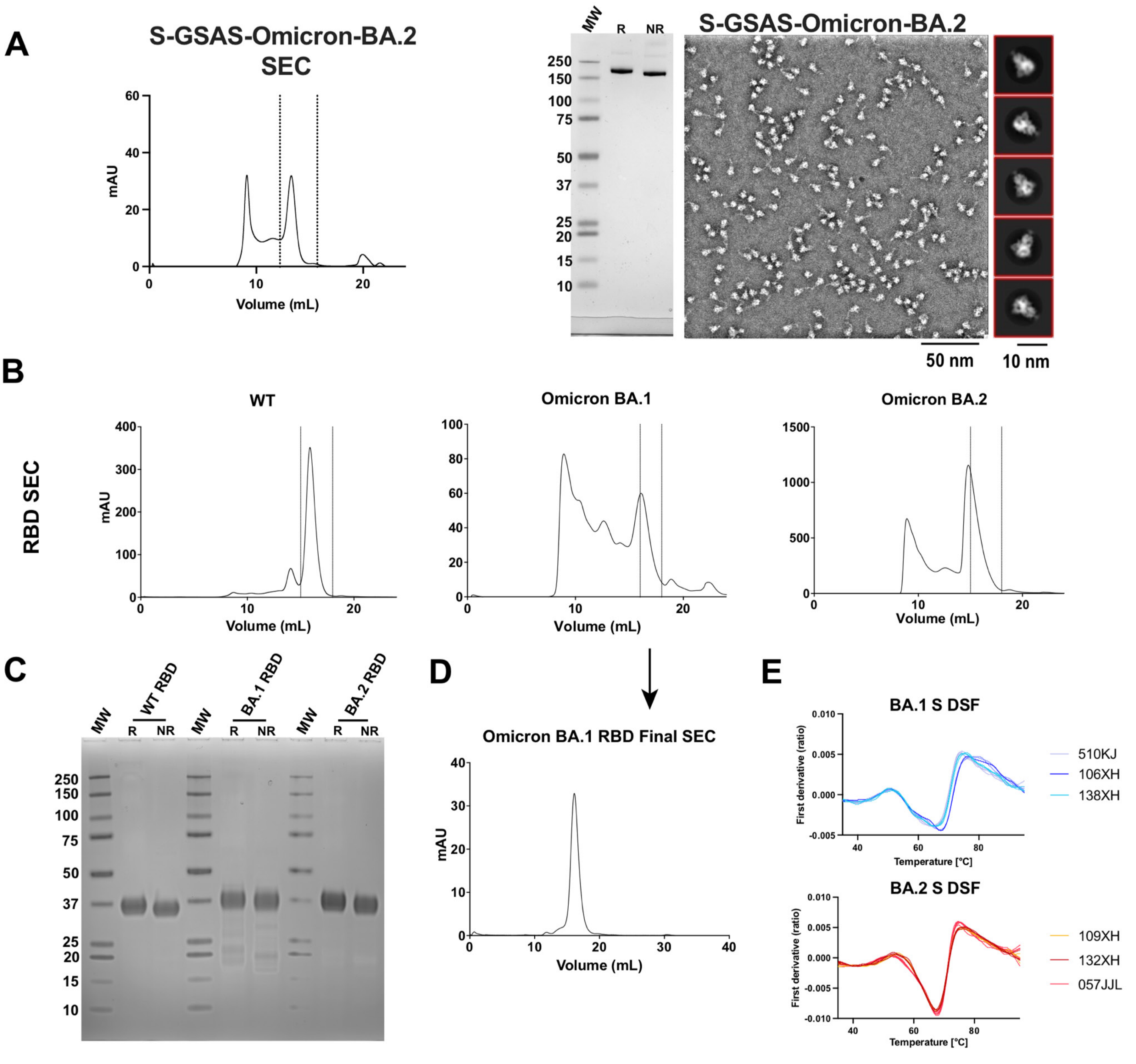


| | Lot # | Ti#1 Mean ± St Dev | Ti#2 Mean ± St Dev | Ti#3 Mean ± St Dev |
|---|---|---|---|---|
| S-GSAS-Omicron-BA.1 | 510KJ | 50.7 ± 0.1 | 64.9 ± 0.9 | 74.1 ± 0.3 |
| | 106XH | 50.8 ± 0.2 | 67.4 ± 0.2 | 77.0 ± 0.9 |
| | 138XH | 50.9 ± 0.9 | 65.4 ± 0.5 | 75.2 ± 0.6 |
| S-GSAS-Omicron-BA.2 | 109XH | 54.5 ± 0.4 | 67.5 ± 0.1 | 76.0 ± 0.8 |
| | 132XH | 53.6 ± 0.6 | 67.4 ± 0.3 | 76.3 ± 0.5 |
| | 057JJL | 52.5 ± 1.0 | 67.6 ± 0.2 | 74.9 ± 0.3 |

**Fig S2. Quality control of SARS-CoV-2 S proteins and RBD proteins. Related to Figures 1 and 2. (A)** S-GSAS-Omicron-BA.2: (left) Size Exclusion Chromatography (SEC) with dashed lines indicating peak fractions pooled for final sample. (middle-left) SDS-PAGE: lane 1, molecular weight marker, lane 2 Reduced (R) and Non-reduced (NR) samples of 2µg Spike protein. (middle-right) Representative micrograph with 50nm scale bar, and (right) 2D class averages with 10nm scale bar. **(B)** From left to right, SEC curves of WT, Omicron-BA.1, and Omicron-BA.2 RBDs, with dashed lines showing pooled fractions. **(C)** RBD protein SDS-PAGE with molecular weight marker, WT R and NR, Omicron-BA.1 R and NR, and Omicron-BA.2 R and NR. **(D)** Second and final SEC of Omicron-BA.1 RBD taken from pooled fractions indicated in B. **(E)** Independently determined DSF plots of S-GSAS-Omicron BA.1 and BA.2 from different preparations of the S proteins. Measurements for each lots included three technical replicates, except 057JJL which had 6. Bottom table shows inflection temperatures ± standard deviation. *Fig 2A also includes data from 106XH and 109XH.

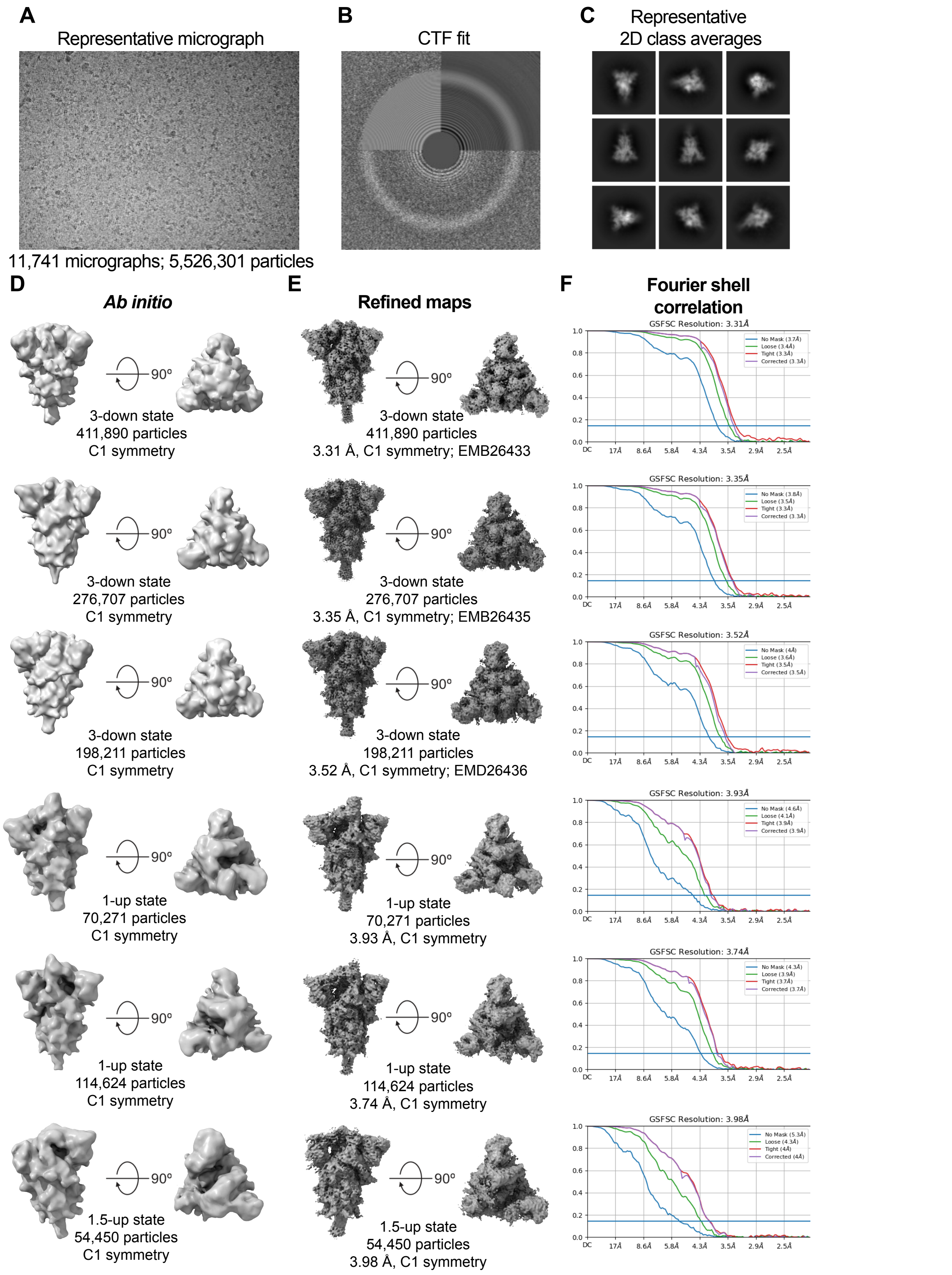


**Fig S3. Cryo-EM data processing for the S-GSAS-Omicron BA.2 ectodomain, Related to Figures 1, 3, 4, 6 and 7, Table 1. (A)** Representative micrograph. **(B)** CTF Fit. **(C)** Representative 2D class averages from cryo-EM dataset. **(D)** Ab initio reconstructions for the cryo-EM 3-RBD-down, 1-RBD-up, and 1.5-RBD-up states. **(E)** Refined cryo-EM reconstructions for the corresponding states. **(F)** Fourier shell correlation curves for the corresponding states.

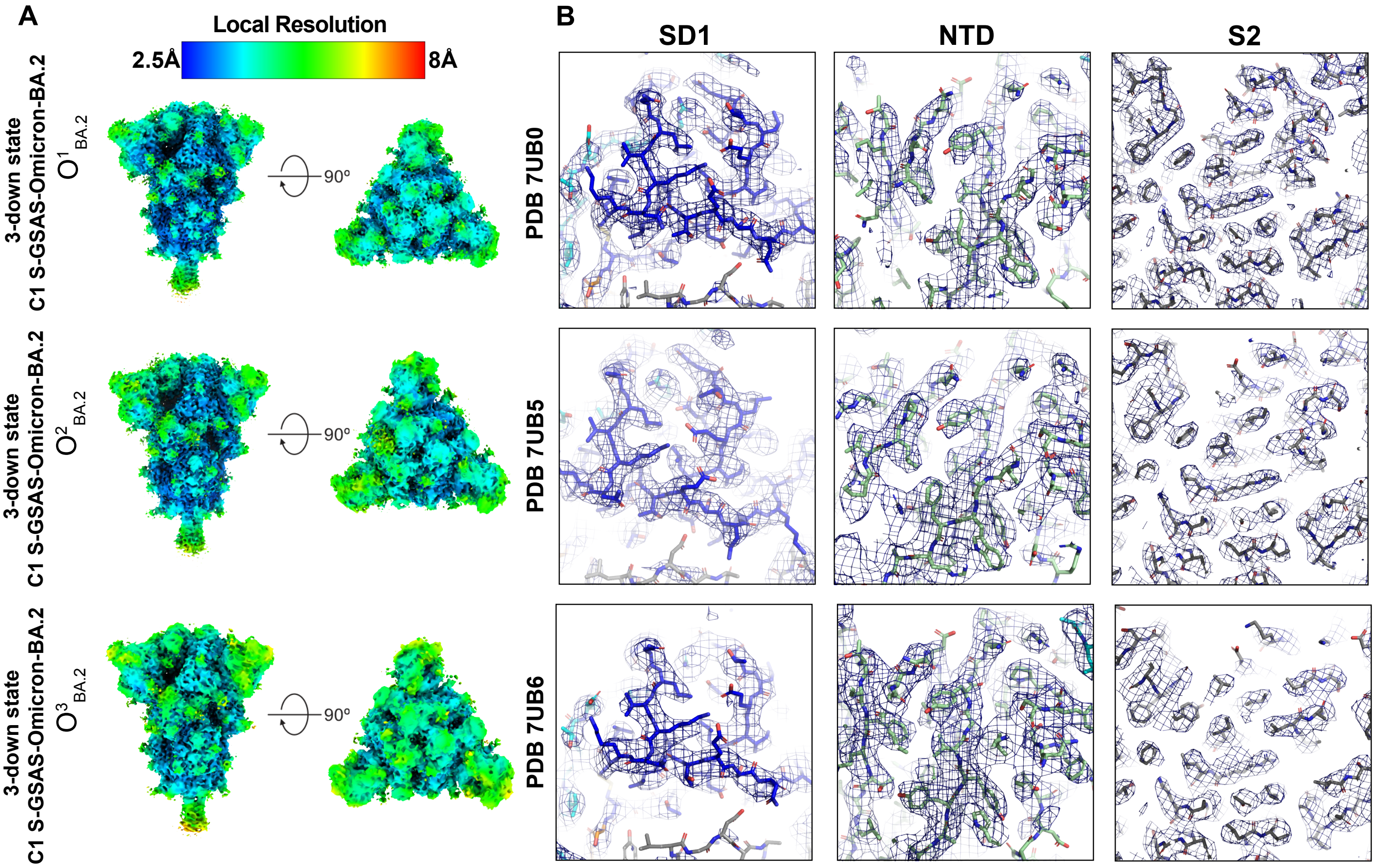


**Fig S4. Quality assessment of the cryo-EM map and model fitting, Related to Figures 1, 3, 4, 6 and 7, Table 1. (A)** Refined maps colored by local resolution ranging from 2.5Å to 8Å. **(B)** Zoomed in views of SD1, NTD, and S2.

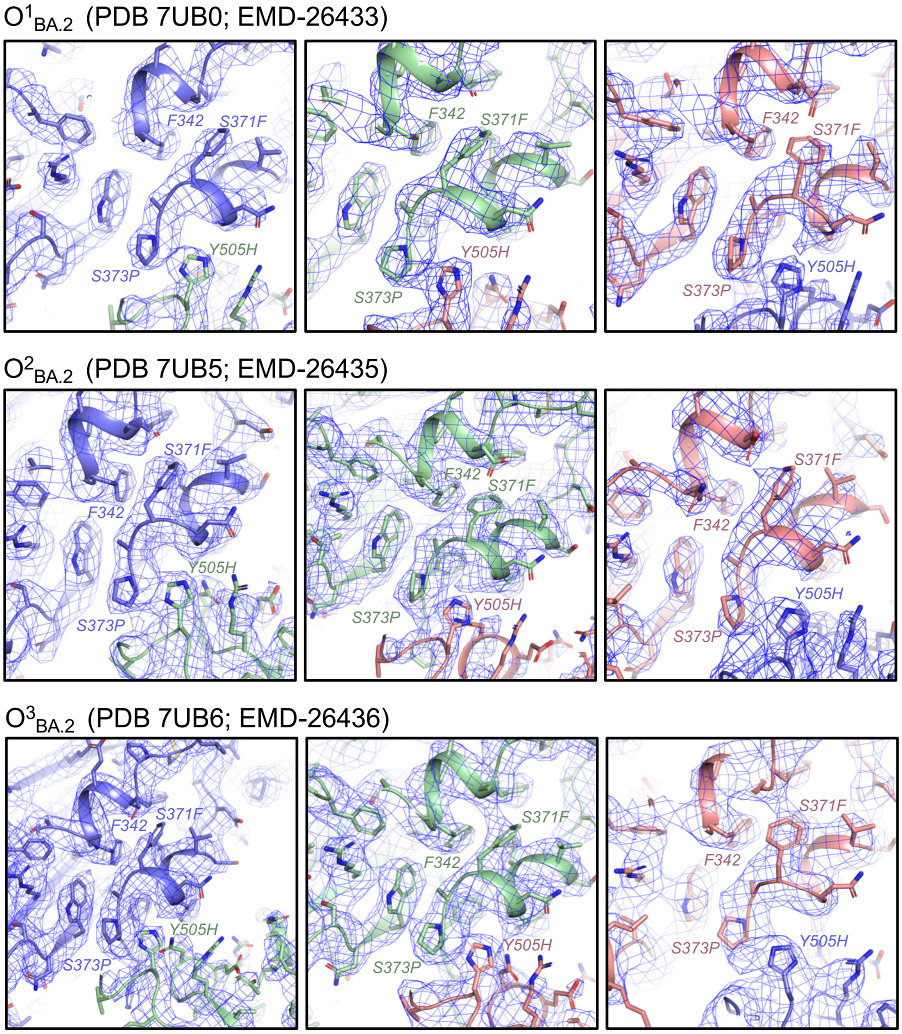


**Fig S5. View of RBD interfacial region bearing S371F and S373 substitutions. Related to Figures 1, 3 and 6.** Cryo-EM reconstructions are shown as blue mesh with underlying fitted model in cartoon and side chains in sticks. Each panel shows the packing of the RBD region bearing the S371F and S373P substitutions with the RBD helix bearing residue F342, as well as inter-protomer packing with the adjacent RBD Y505H substitution stacking against P373. Details are shown for each protomer in $O^1_{BA.2}$ (top row), $O^2_{BA.2}$ (middle row) and $O^3_{BA.2}$ (bottom row).

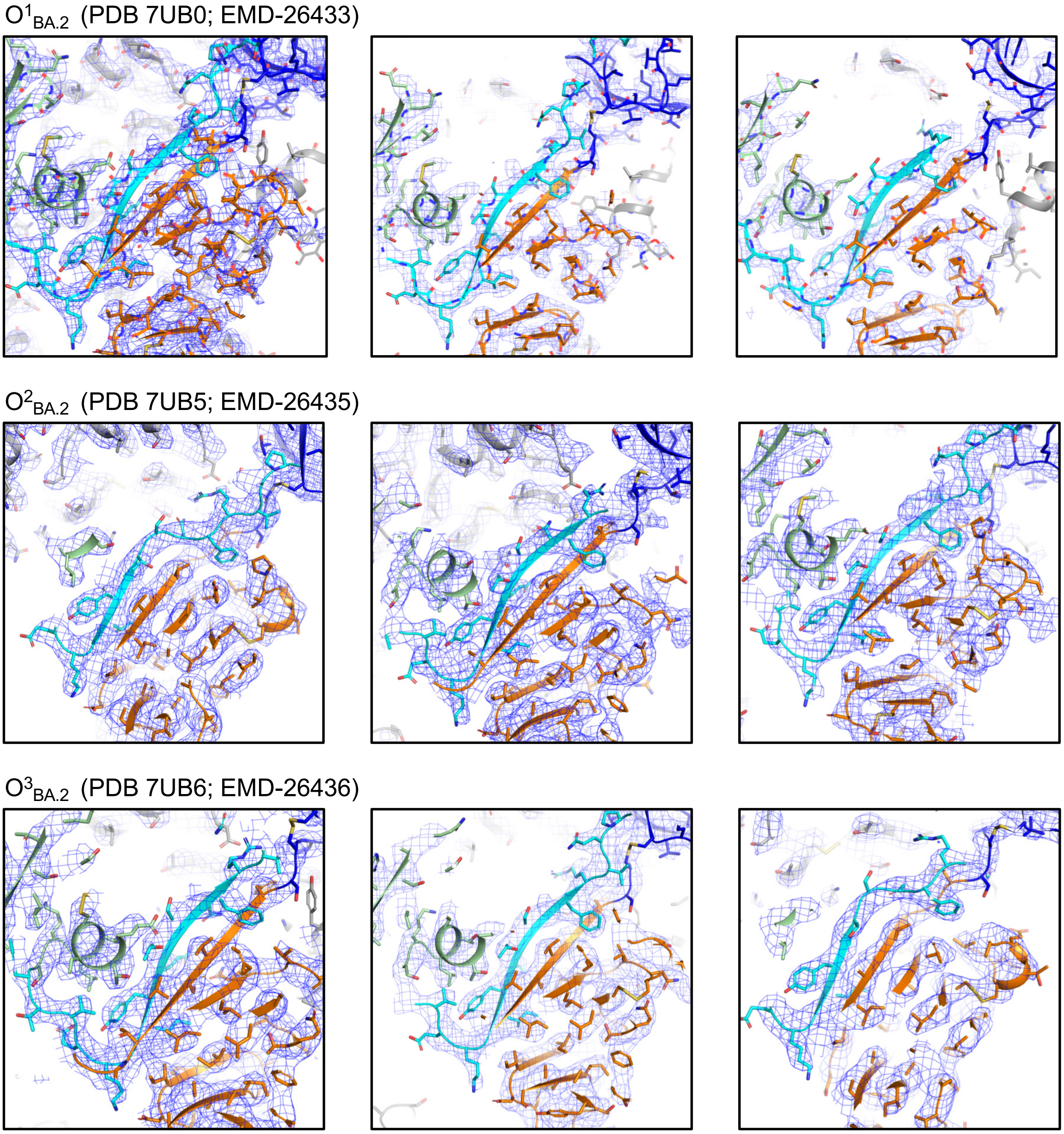


**Fig S6. View of NTD-to-RBD (N2R) linker in 3-RBD-down Omicron BA.2 S. Related to Figures 1 and 4.** Cryo-EM reconstructions are shown as blue mesh with underlying fitted model in cartoon and side chains in sticks. Each panel shows the N2R linker (cyan) of a protomer stacked against its SD2 subdomain (orange). The NTD is colored green and SD1 subdomain blue. Details are shown for each protomer in $O^1_{BA.2}$ (top row), $O^2_{BA.2}$ (middle row) and $O^3_{BA.2}$ (bottom row).

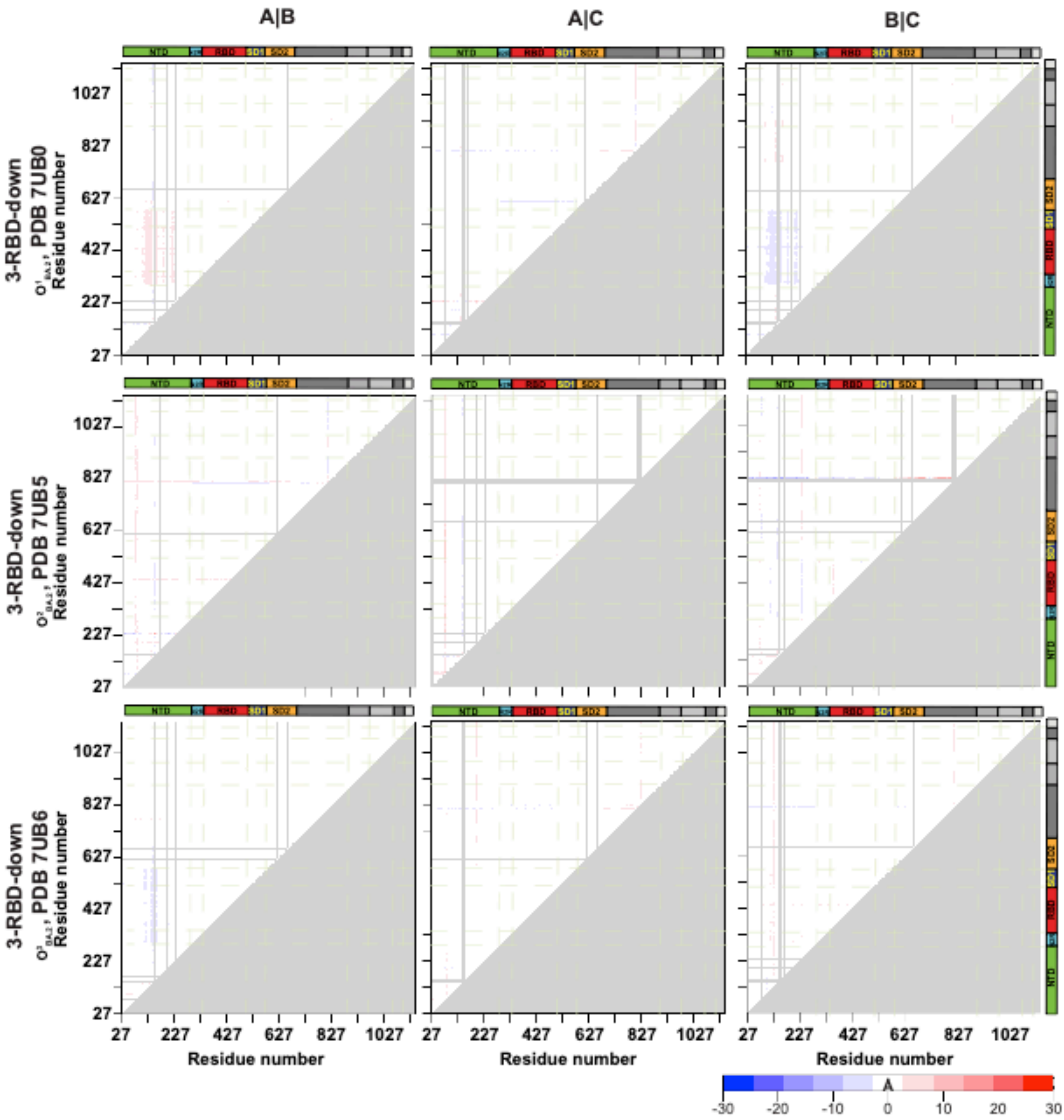


**Figure S7. Intrachain variability in the S-GSAS-Omicron-BA.2 S ectodomain 3-RBD-down structures. Related to Figure 1.** Difference distance matrices (DDMs) provide superposition-free comparisons between a pair of structures by calculating the differences between the distances of each pair of Ca atoms in a structure and the corresponding pair of Ca atoms in the second structure. DDM of the 3-RBD down structures comparing each chain of a structure to the other chains of the same structure. The coloring scale used for the DDM analysis is represented on the bottom right corner.

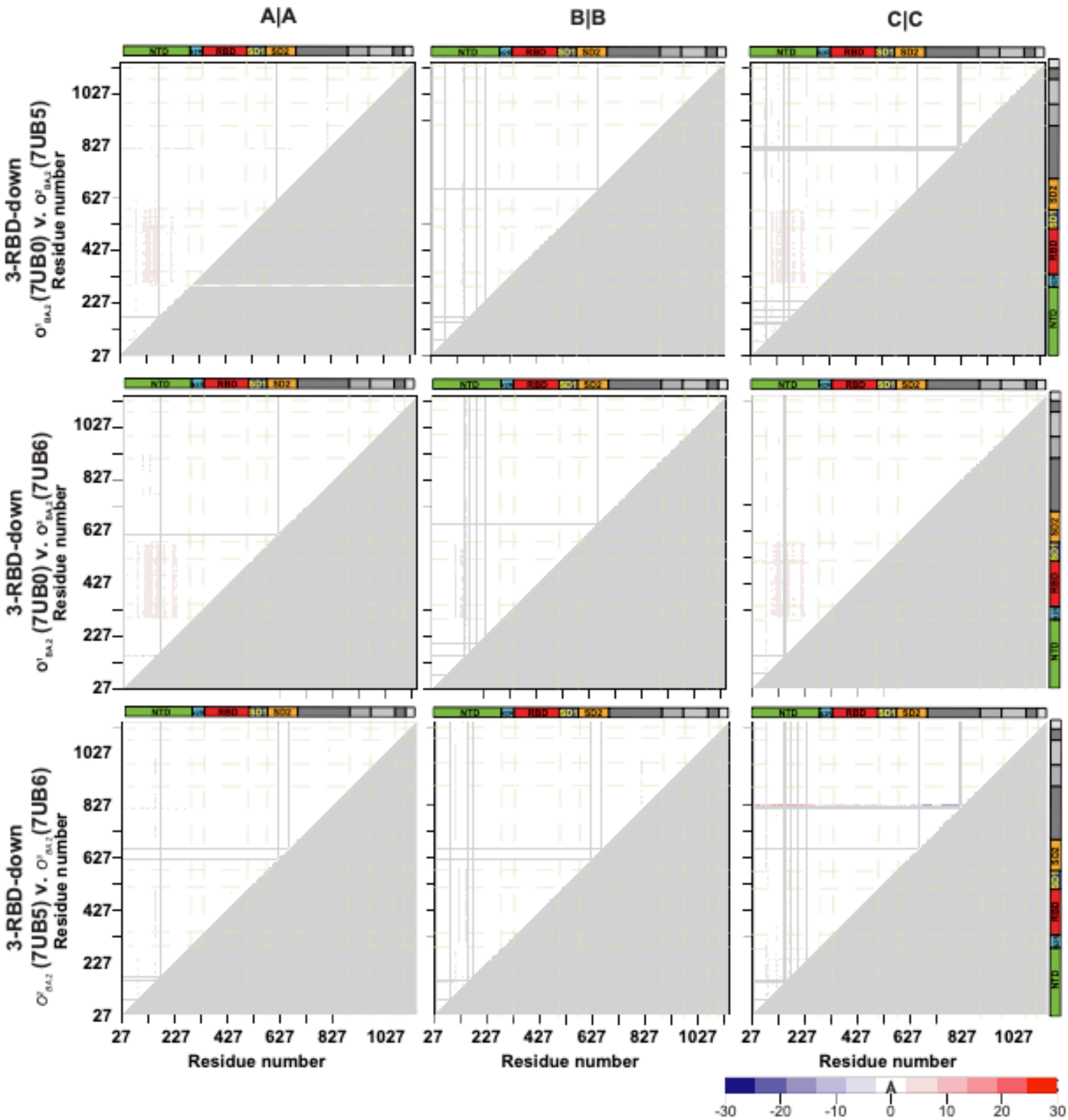


**Figure S8. Interchain variability in the S-GSAS-Omicron-BA.2 S ectodomain 3-RBD-down structures. Related to Figure 1.** DDM of the 3-RBD down structures comparing each chain of a structure to the same chain of another structure. The coloring scale used for the DDM analysis is represented on the bottom right corner.

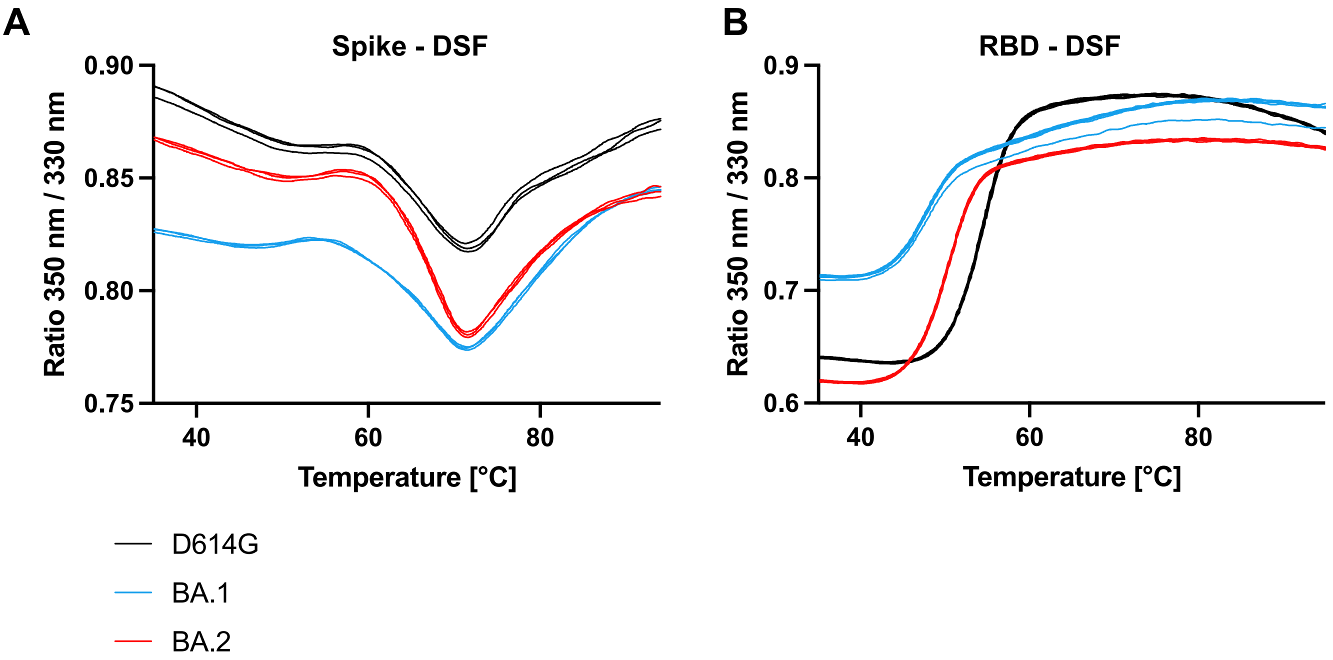


**Fig S9. Thermostability of SARS-CoV-2 S ectodomain and monomeric RBD proteins. Related to Figure 2.** DSF profiles of **(A)** SARS-CoV-2 S ectodomain and **(B)** monomeric RBD constructs, showing changes in protein intrinsic fluorescence (expressed as a ratio between fluorescence at 350 and 330 nm). For each S protein construct (color coded as shown), three overlaid curves (technical replicates) are shown.

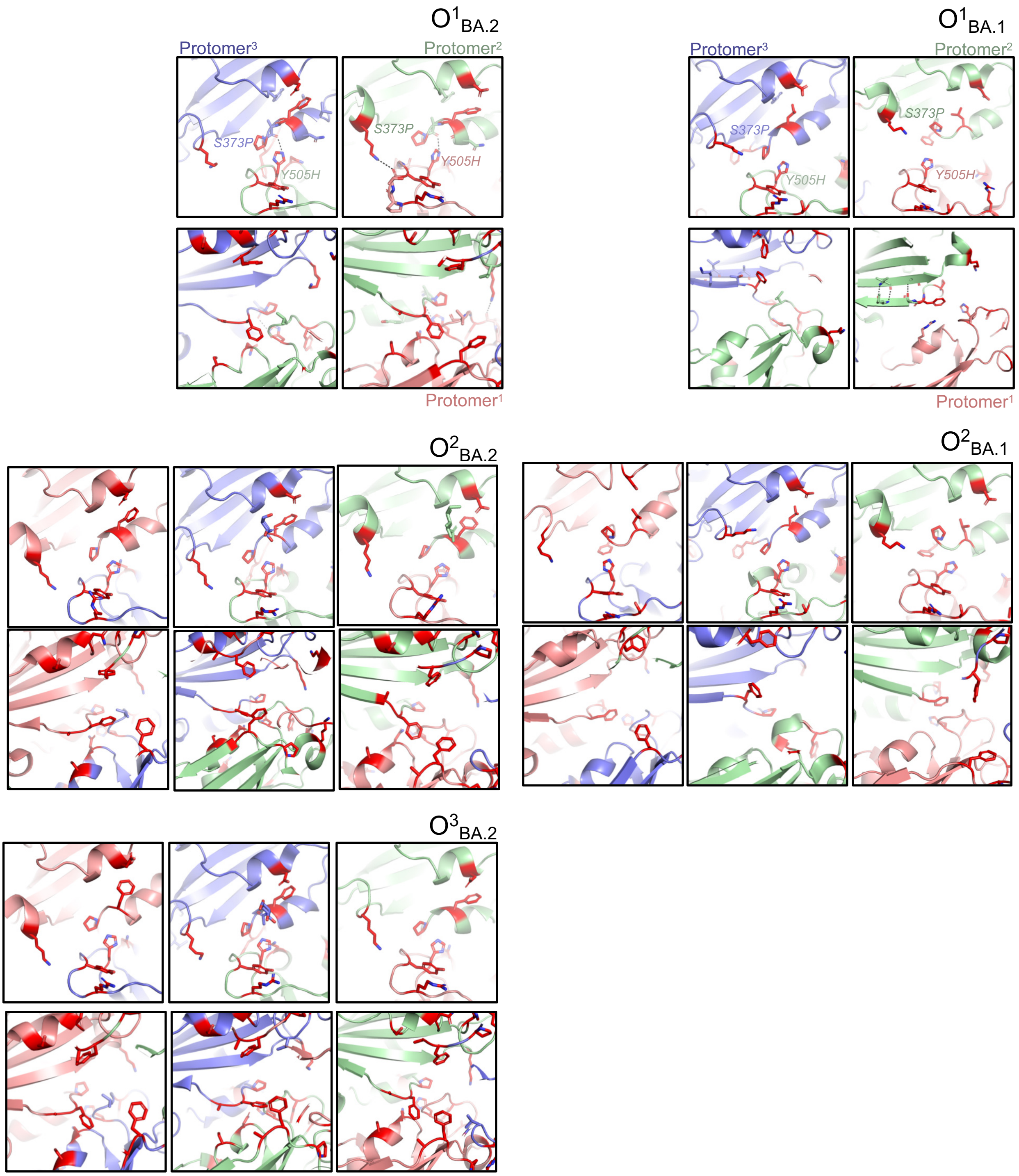


**Fig S10. Comparison of the RBD-RBD interface in the Omicron BA.1 and BA.2 S 3-RBD-down structures, Related to Figure 3.** Interprotomer interactions between adjacent RBDs are shown for (left) the BA.2 S protein structures ($O^1_{BA.2}$: PDB 7UB0; $O^2_{BA.2}$: PDB 7UB5; $O^3_{BA.2}$: PDB 7UB6), and (right) the BA.1 S protein structures ($O^1_{BA.1}$: PDB 7TF8; $O^2_{BA.1}$: PDB 7TL1)

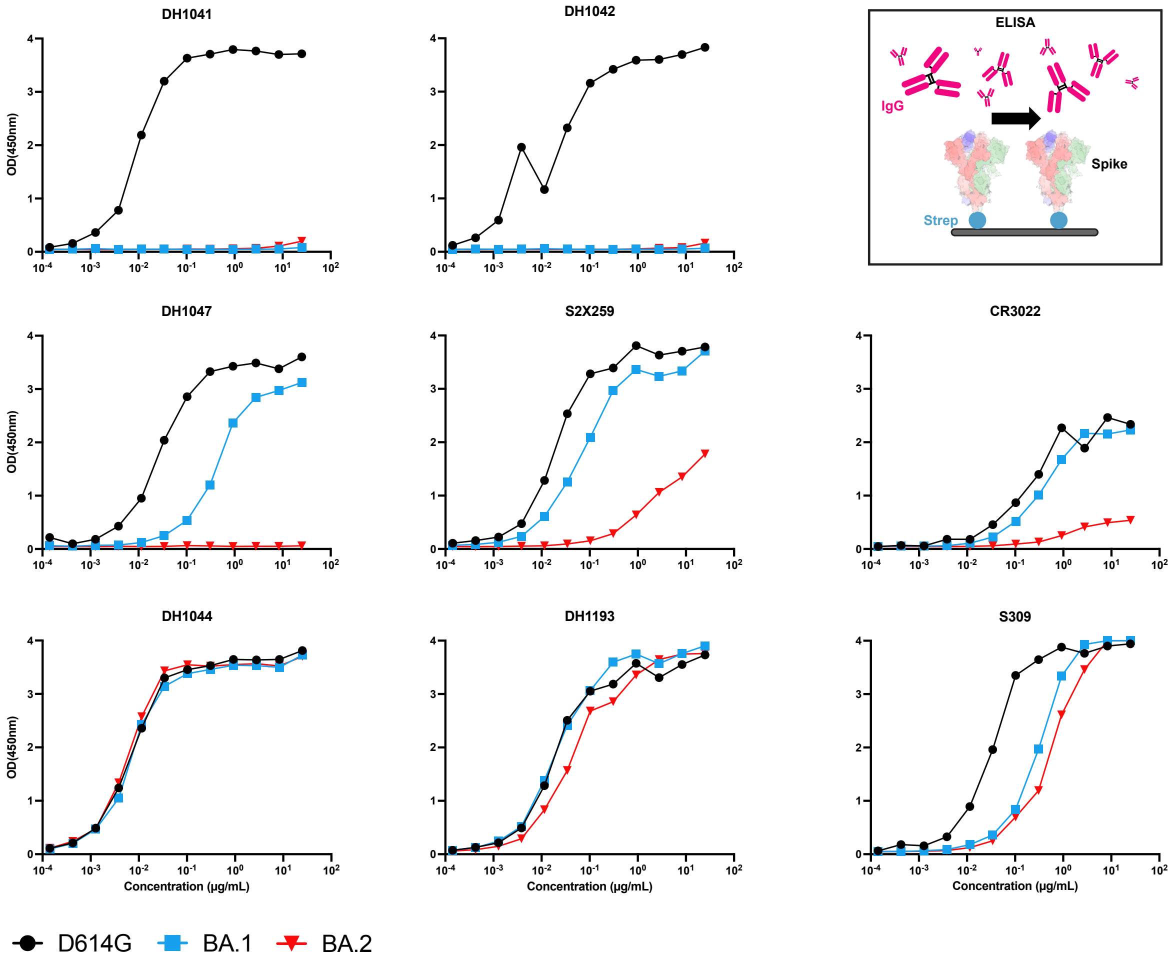


**Fig S11. Binding of RBD-directed antibodies to spike variants measured by ELISA, Related to Figures 5 and 6.** ELISA binding of antibodies, top row, DH1041 and DH1042 (RBM-targeting), second row, DH1047, S2X259, and CR3022 (inner face RBD-targeting), and bottom row, DH1044, DH1193, and S309 (outer face RBD-targeting) to the D614G (black), BA.1 (blue) and BA.2 (red) S protein ectodomains. Schematic shows the assay format.

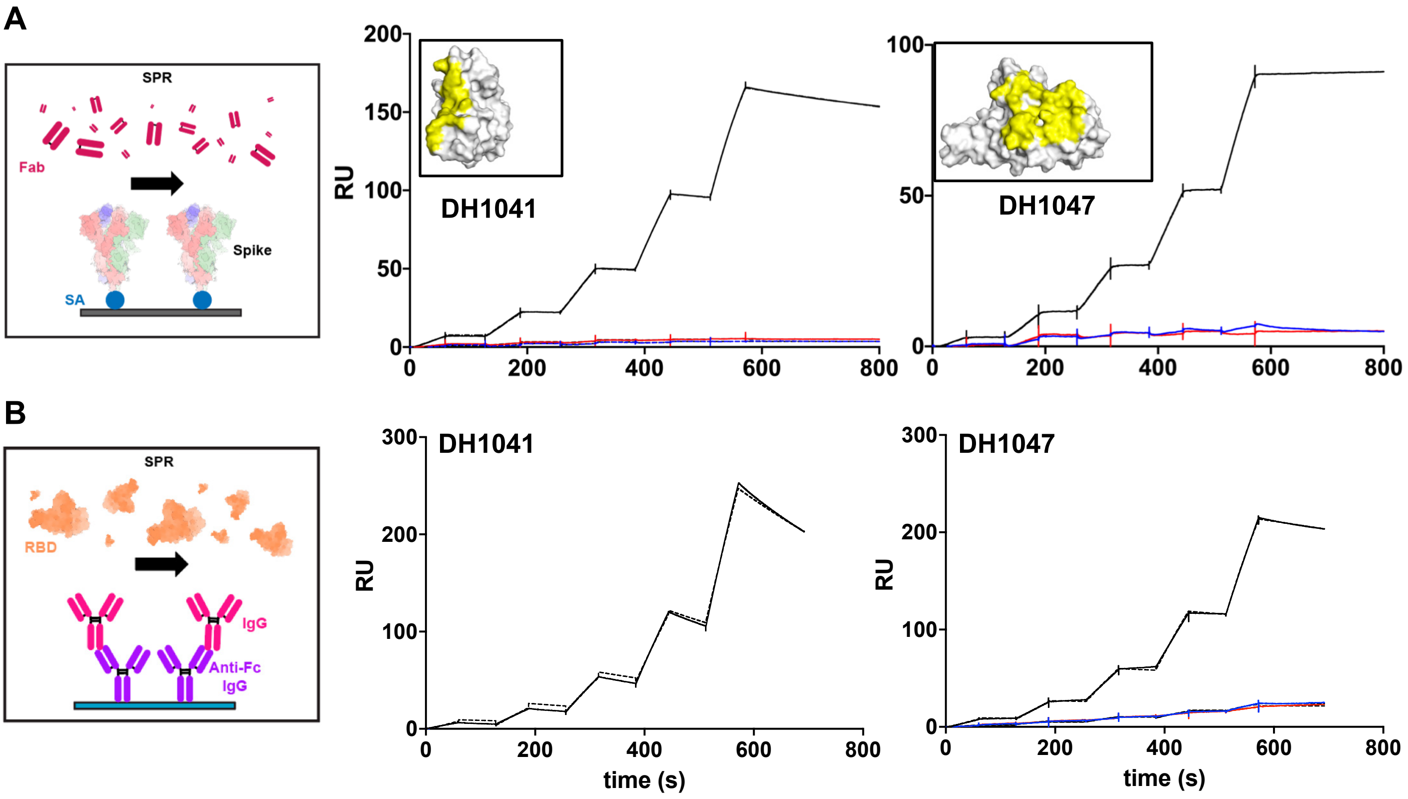


**Fig S12. Binding of RBD-directed Fabs to S protein and RBD variants measured by SPR, Related to Figures 5 and 6. A.** Binding of RBD-directed antibodies DH1041 and DH1047 to the D614G (black), BA.1 (blue) and BA.2 (red) S protein ectodomains measured by SPR. Schematic shows the assay format. **B.** Binding of RBD-directed antibodies DH1041 and DH1047 to the WT (black), BA.1 (blue) and BA.2 (red) monomeric RBD constructs measured by SPR. Schematic shows the assay format. The solid lines are the binding sensograms; the dotted lines show fits of the data to a 1:1 Langmuir binding model.

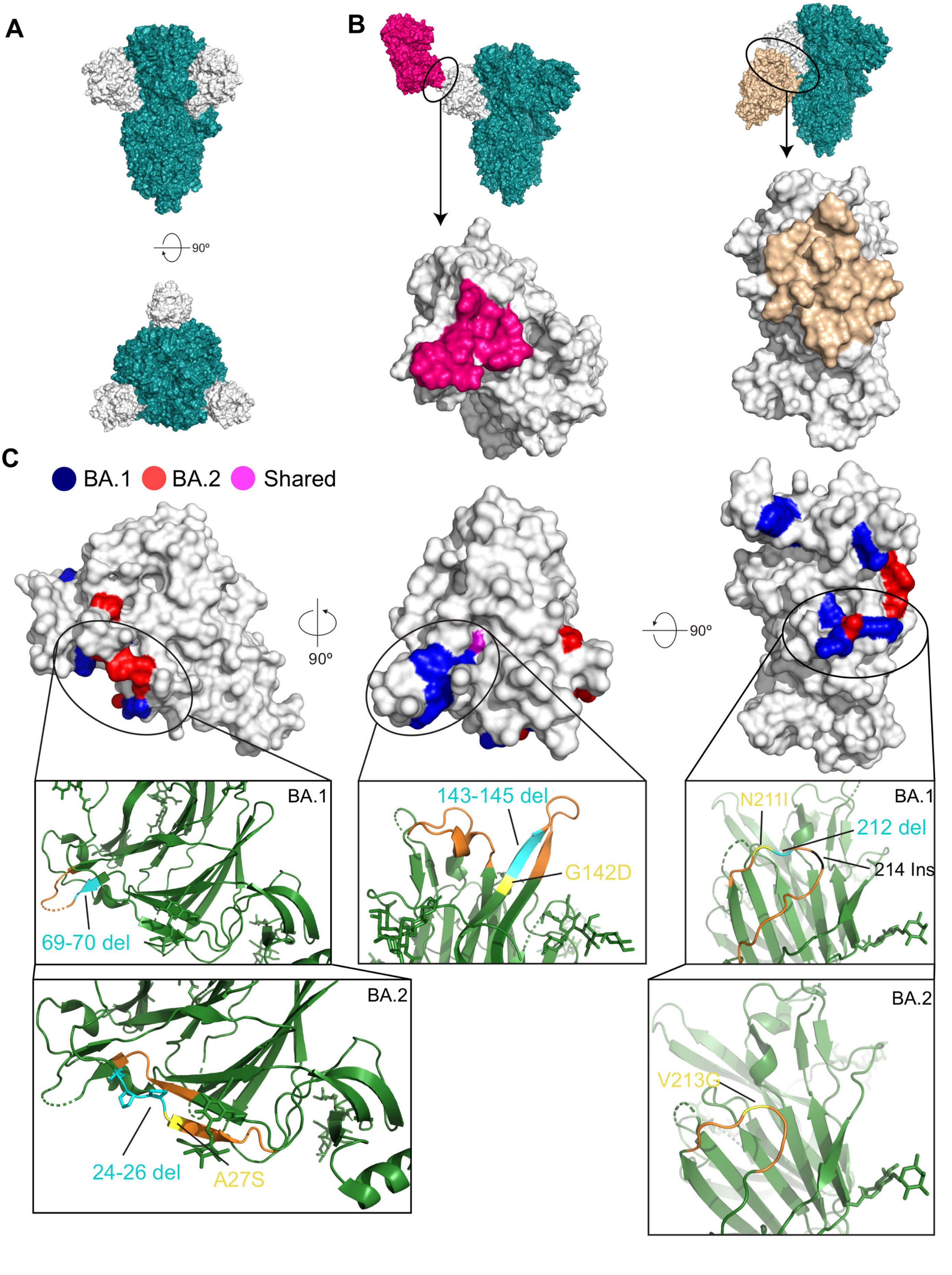
A
90º
B
C
BA.1
BA.2
Shared
90º
90º
BA.1
69-70 del
143-145 del
G142D
N211I
212 del
214 Ins
BA.2
24-26 del
A27S
V213G

**Fig S13. Comparison of BA.1 and BA.2 N-terminal domain (NTD) mutations. Related to Figure 5. (A)** Surface representation of the SARS-CoV-2 S ectodomain, colored teal, with each NTD colored white, side view and top view. (**B)** (Top) Surface views of the spike, with Fab regions of NTD-directed antibodies bound: DH1050.1, pink, and DH1052, gold. Only one NTD is colored white and shown bound to the Fab. (Bottom) One NTD shown in surface representation with epitopes of the NTD-directed antibodies shown in the same colors as the antibodies in the top panel. **(C)** Three views of the wild-type NTD with residues mutated in the Omicron variants highlighted as follows: blue: BA.1, red: BA.2, magenta: shared in BA.1 and BA.2. Rectangular panels show zoomed-in views in cartoon representation of areas highlighted in C, with mutated residues shown as sticks, with the following color scheme: yellow: mutated residues, cyan: deleted residues, black: insertion site, orange: nearby residues/loop residues, other NTD residues: green. Models used: A: 7LAB, B: 7LCN (DH1050.1-bound SARS-CoV-2 2P-S ectodomain) and 7LAB (DH1052-bound SARS-CoV-2 2P-S ectodomain), C: 7LY3

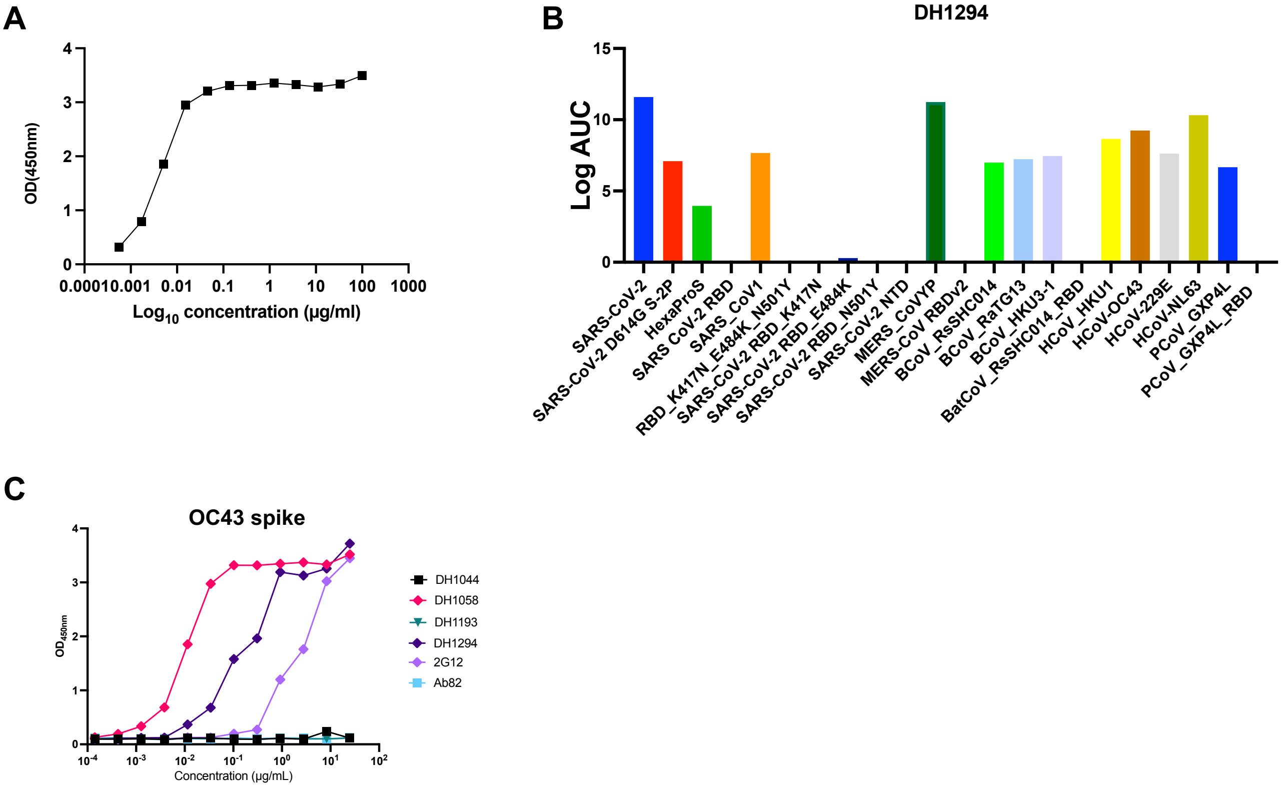


**Figure S14. Epitope mapping of fusion peptide directed antibody DH1294. Related to Figure 7. (A)** Binding of antibody DH1294 to the SARS-CoV-2 fusion peptide (the sequence of the peptide used in the ELISA is shown in Figure 7H)**. B.** ELISA binding of DH1294 to diverse CoV spikes. **C.** ELISA binding of antibodies to the CoV OC43 spike.

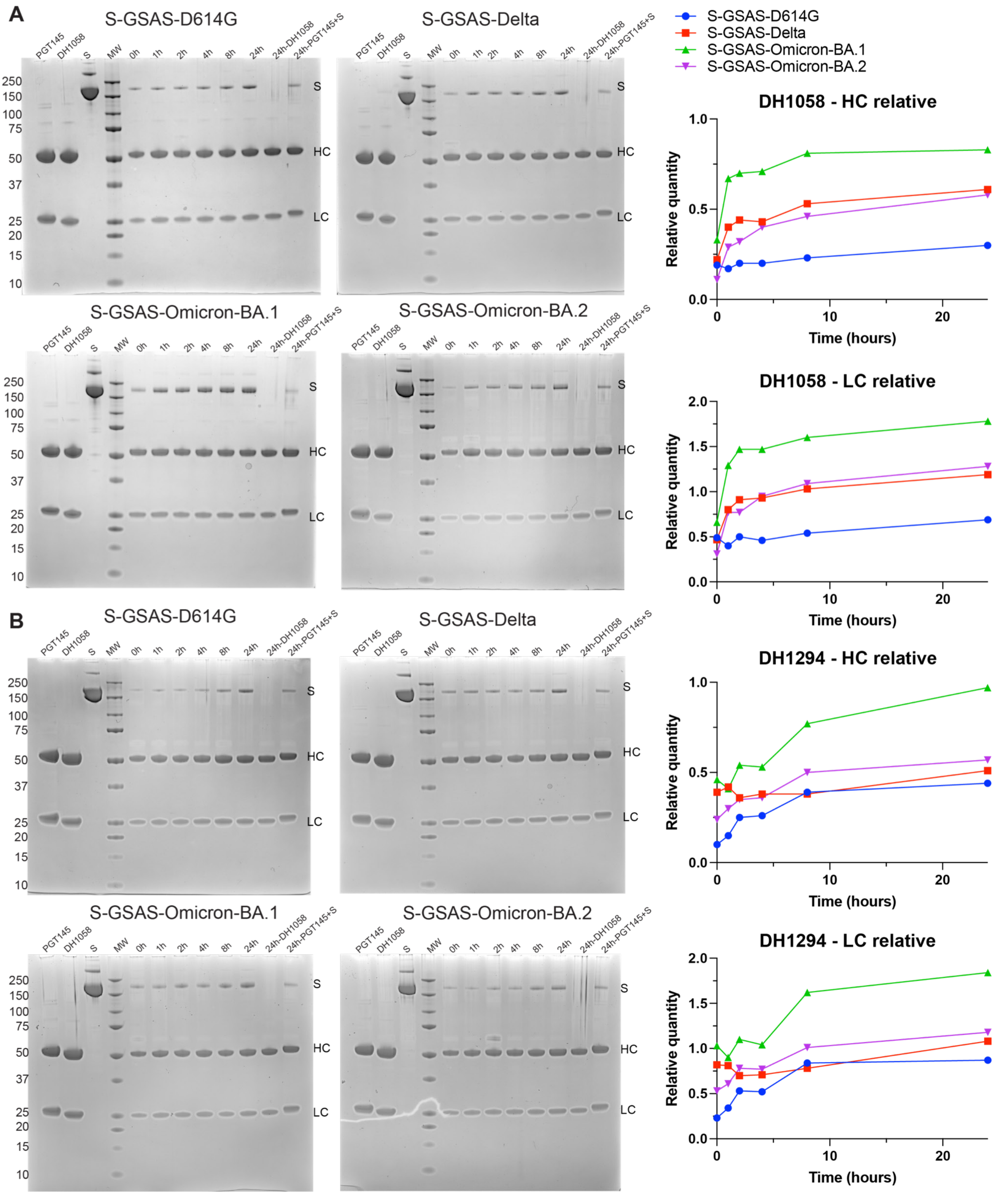

A
S-GSAS-D614G
S-GSAS-Delta
S-GSAS-Omicron-BA.1
S-GSAS-Omicron-BA.2
PGT145
DH1058
S
MW
0h
1h
2h
4h
8h
24h
24h-DH1058
24h-PGT145+S
250
150
100
75
50
37
25
20
15
10
HC
LC
B
S-GSAS-D614G
S-GSAS-Delta
S-GSAS-Omicron-BA.1
S-GSAS-Omicron-BA.2
DH1058 - HC relative
DH1058 - LC relative
DH1294 - HC relative
DH1294 - LC relative
Relative quantity
Time (hours)
0.0
0.5
1.0
1.5
2.0
0
10
20

**Fig S15. Time dependent exposure of fusion peptide (FP) to FP-directed antibodies. Related to Figure 7. A.** Left. Reducing SDS-PAGE of immunoprecipitation of S-GSAS-D614G, S-GSAS-Delta, S-GSAS-Omicron-BA.1 and S-GSAS-Omicron-BA.2 using FP-directed antibody DH1058 bound to a Protein A coupled resin. Right. Intensity of the S protein band at each time point was normalized with the intensity of the heavy chain (HC, top) or light chain (LC, bottom) band. **B.** Same as **A.** but for antibody DH1294.

| | D614G | | Omicron-BA.1 | | Omicron-BA.2 | | Omicron-BA.3 | |
|---|---|---|---|---|---|---|---|---|
| | ID50 | ID80 | ID50 | ID80 | ID50 | ID80 | ID50 | ID80 |
| DH1044 | 0.04 | 0.17 | 0.52 | 2.3 | 0.09 | 0.19 | 0.35 | 0.99 |
| DH1047 | 0.1 | 0.34 | >25 | >25 | >25 | >25 | >25 | >25 |
| DH1193 | 1 | 3.7 | 2 | 11 | 6.1 | 16 | 1.3 | 4.4 |

**Table S1. Pseudovirus neutralization by RBD-directed antibodies DH1044, DH1047 and DH1193.**
